# Brain connectivity at rest predicts individual differences in normative activity during movie watching

**DOI:** 10.1101/2021.01.06.425410

**Authors:** David C. Gruskin, Gaurav H. Patel

**Author notes:** Corresponding author **Correspondence:** Gaurav H. Patel, 1051 Riverside Drive Unit 21, New York, NY 10032. **Author email addresses:**.

## Abstract

When multiple individuals are exposed to the same sensory event, some are bound to have less typical experiences than others. These atypical experiences are underpinned by atypical stimulus-evoked brain activity, the extent of which is often indexed by intersubject correlation (ISC). Previous research has attributed individual differences in ISC to variation in trait-like behavioral phenotypes. Here, we extend this line of work by showing that an individual’s degree and spatial distribution of ISC are closely related to their brain’s intrinsic functional architecture. Using resting state and movie watching fMRI data from 176 Human Connectome Project participants, we reveal that resting state functional connectivity (RSFC) profiles can be used to predict cortex-wide ISC with considerable accuracy. Similar region-level analyses demonstrate that the amount of ISC a brain region exhibits during movie watching is associated with its connectivity to others at rest, and that the nature of these connectivity-activity relationships varies as a function of the region’s role in sensory information processing. Finally, we show that an individual’s unique spatial distribution of ISC, independent of its magnitude, is also related to their RSFC profile. These findings suggest that the brain’s ability to process complex sensory information is tightly linked to its baseline functional organization and motivate a more comprehensive understanding of individual responses to naturalistic stimuli.

## 1. INTRODUCTION

The ability to interpret information from the outside world in a manner similar to one’s conspecifics is critical to healthy human behavior. Recent work on the neural substrates of sensory information processing has accordingly focused on the processing of naturalistic stimuli that reflect the complex and continuous nature of everyday experiences. One key finding from this line of research is that rich, time-locked stimuli such as movies elicit strongly conserved brain responses across individuals (Hasson et al., 2004). The extent of this stimulus-evoked brain activity is most frequently indexed by intersubject correlation (ISC), a data-driven measure of how similar one’s activity time course in a given brain region is to that of another individual or group average (Hasson et al., 2004; Nastase et al., 2019).

Early research using naturalistic stimuli focused on aspects of stimulus-evoked activity that are shared across subjects, but individual differences in ISC have become a target of recent work (Gruskin et al. 2020; Salmi et al. 2013; Campbell et al. 2015; Finn et al. 2018; Finn et al. 2020; Guo et al. 2015). These studies have identified several cognitive and clinical correlates of between-subject variance in ISC, where a low value for a given individual reflects atypical processing of the stimulus relative to the rest of the sample. Specifically, this measure of functional typicality has been associated with trait-like phenotypes (e.g., attentional control, Campbell et al., 2015) in healthy subjects and symptom severity (e.g., depression questionnaire scores, Gruskin et al., 2020; Guo et al., 2015) in clinical/sub-clinical populations. The aim of the present manuscript is to explore a different, but not mutually exclusive hypothesis; namely, that the degree to which an individual exhibits normative activity during movie watching is related to their brain’s intrinsic functional architecture.

Just as brain responses to naturalistic stimuli reflect individual-specific variations on a population-general theme, the brain’s resting functional architecture is organized into modular networks whose general structure is conserved from person to person but whose fine-grained topologies and strengths serve as stable identifiers of individuals (Finn et al., 2015; Gordon et al., 2017; van den Heuvel & Hulshoff Pol, 2010). These networks are classically derived from resting state functional connectivity (RSFC), which quantifies the statistical dependence between activity time courses from any two regions measured while the brain is not performing an explicit task. A growing literature has shown that RSFC can be used to predict both normal and pathological variation in model-based measures of brain activity during cognitively demanding tasks, providing compelling evidence for the idea that RSFC describes the routes along which task-relevant information travels (Cole et al., 2016; Ito et al., 2017; Mill et al., 2020; Tavor et al., 2016; van den Heuvel & Hulshoff Pol, 2010). Despite these recent advances, the extent to which RSFC can (1) account for individual variability in model-free neural responses to naturalistic stimuli and (2) contribute to our understanding of how the brain processes such stimuli is still unclear.

To characterize how resting state functional connectivity is associated with normative stimulus-evoked activity during movie watching, we analyzed fMRI data acquired across multiple days from healthy Human Connectome Project participants. First, we demonstrate that models trained on resting state and movie watching data can predict ISC from RSFC in held-out subjects and in data from held-out stimuli. Next, we show that the resting state functional connections most associated with ISC in a specific region vary systematically across the brain and quantify how greater intrinsic connectivity to one region is associated with more or less typical activity in others. Finally, we show that individual-specific RSFC patterns are related to the spatial distribution of ISC across cortex. Taken together, these results suggest that the brain’s intrinsic functional architecture is closely related to its ability to process audiovisual information, providing both important context for interpreting atypical responses to naturalistic stimuli and new evidence for the neural underpinnings of sensory information processing.

## 2. METHODS

### 2.1 Participants

Data used for this project come from the Human Connectome Project (HCP) Young Adult 7T release (Van Essen et al., 2013). Of the 184 subjects who underwent 7T fMRI scanning, eight participants did not complete every resting state and movie watching run. These subjects were excluded from all analyses for a sample size of n = 176 (106 females, 70 males). All participants were healthy individuals between the ages of 22 and 36 (mean age = 29.4 years, standard deviation = 3.3). Self-reported racial identity in this sample was 87.5% White, 7.4% Black or African American, 4% Asian/Native Hawaiian/Other Pacific Islander, and 1% unknown/not reported, and 1.7% of the sample identified as Hispanic/Latino.

### 2.2 fMRI data

FMRI data for the HCP 7T release were collected at the University of Minnesota on a 7T Siemens Magnetom scanner during four sessions spread across two or three days. Each day of data collection involved two resting state and two movie watching runs. The same echo-planar imaging sequence was used for all rest and movie scans and its key parameters are as follows: time of repetition (TR) = 1000 ms, echo time (TE) = 22.2 ms, number of slices = 85, flip angle = 45 degrees, spatial resolution = 1.6 mm^3^.

All four rest runs had a duration of 900 TRs/15:00 minutes, whereas the four movie runs had variable durations (921, 918, 915, and 901 TRs). Subjects were instructed to fixate on a projected bright crosshair on a dark background during the rest runs and passively viewed a series of video clips (with sound) during each movie run. Movie runs one and three each consisted of four unique clips from independent films, and movie runs two and four were each composed of three unique clips from major motion pictures. The same montage of brief (1-4 seconds) videos was included at the end of every run for test-retest purposes. Clip durations varied between 64 and 259 seconds. More information on these clips can be found in Finn & Bandettini (2020) and at https://db.humanconnectome.org. Each video clip was preceded by 20 seconds of rest (during which the word “REST” was projected against a dark background). Any TR that took place during one of these rest blocks was discarded from all analyses. The first 20 TRs after each rest block (corresponding to the start of each video clip) were also discarded to prevent onset transients from biasing our intersubject correlation measurements (Dosenbach et al., 2006; Fox et al., 2005), leaving four movie runs of 722, 755, 716, and 738 TRs, respectively. Finally, rest and movie runs from the same day were normalized and concatenated, yielding one rest run (1800 TRs) and one movie run (1477/1454 TRs) for each of the two days of data collection.

### 2.3 Preprocessing and parcellation

ICA-FIX denoised CIFTI files (e.g., *rfMRI_REST1_7T_PA_Atlas_hp2000_clean*.*dtseries*.*nii*) at 2 mm resolution were downloaded from ConnectomeDB. Briefly, these data were pre-processed using motion and distortion correction, high-pass temporal filtering, and MNI alignment, followed by regression of 24 motion parameters as well as a set of independent component analysis-derived confound time courses (Glasser et al., 2013; Griffanti et al., 2014). Because head motion-related and other artifacts may persist in fMRI data even after ICA-FIX, additional denoising was performed. Following recent work with HCP data (Finn & Bandettini, 2020; Li et al., 2019), the average time course of all grayordinates and its temporal derivative was regressed from each scan. Additionally, high-motion frames were censored from all rest scans. To identify these frames, a 0.2 Hz (12 breaths per minute) low pass filter was first applied to each scan’s framewise displacement (FD) trace to account for respiratory artifacts found in fMRI data (Fair et al., 2020; Gratton et al., 2020). Any volume exceeding 0.2 mm FD post-filtering was flagged, as were all runs of fewer than five contiguous volumes. Finally, delta functions corresponding to each of the censored volumes were included along with the two global signal time courses for each rest scan’s denoising design matrix. Frame censoring was not performed on the movie scans to ensure that intersubject correlation would be calculated with the same number of degrees of freedom across individuals.

Following denoising, functional data were parcellated into 360 cortical regions of interest using the Glasser HCP-MMP parcellation (Glasser et al., 2016). The Glasser parcellation was chosen for its anatomically specific labels, association with the HCP dataset and CIFTI format, and compatibility with the Cole-Anticevic Brain-wide Network Partition (Ji et al., 2019), whose granular and interpretable network definitions (e.g., separate language and auditory networks) are especially suitable for the analysis of BOLD signals evoked by complex audiovisual stimuli.

### 2.3 Resting state functional connectivity

Using data from the two concatenated rest scans, time courses from all possible pairs of the 360 parcels were (Pearson) correlated to create two symmetric 360 × 360 resting state functional connectivity (RSFC) matrices for each subject, one for each day of data collection. Frames flagged as having high motion as per section 2.3 were excluded from the correlation calculations.

### 2.4 Intersubject correlation

Intersubject correlation (ISC) analysis was used to quantify the typicality of each individual’s BOLD responses to the Day 1 and Day 2 movie stimuli. For each of the 360 cortical parcels, each participant’s BOLD signal time-course was normalized and (Pearson) correlated with that parcel’s average BOLD signal time course across all other participants to yield a 360 parcels × 176 subjects intersubject correlation matrix. This was repeated separately for both concatenated movie scans such that every participant had two independent ISC values for each of the 360 parcels, each reflecting the typicality of that individual’s BOLD responses to that day’s movie stimuli in that parcel.

### 2.5 Ridge connectome-based predictive modeling

#### Overview

Ridge connectome-based predictive modeling (rCPM) was used to relate RSFC to global ISC (gISC), defined as the average of each participant’s 360 parcel-level ISC values. This measure functions as a simple, cortex-wide indicator of how typical an individual’s brain activity during movie watching is compared to the rest of the group. CPM is an established technique for predicting behavior (e.g., fluid intelligence, symptom severity) from RSFC (Finn et al., 2015; Rosenberg et al., 2016; Shen et al., 2017). Here, we use a variant of CPM, rCPM (Gao et al., 2019), to predict gISC from RSFC. Although the prediction of gISC is conceptually similar to the prediction of any other measure, gISC’s derivation from BOLD data leaves it susceptible to motion and other fMRI acquisition artifacts. More importantly, individual differences in these artifacts are likely consistent across rest and movie scans such that relationships between RSFC and gISC may be driven by artifacts rather than neural signals of interest. Therefore, in addition to the conservative preprocessing approach outlined in section 2.3, all correlations between RSFC and (g)ISC performed as part of the training and testing of the rCPM models as well as elsewhere in the paper included motion estimates (mean FD) as covariates. Although detailed descriptions of CPM can be found elsewhere (Shen et al., 2017), provided below is a summary of the specific approach used in this paper.

#### Generating the cross-validated models

Because the HCP 7T dataset is composed of data from individuals of varying degrees of genetic relatedness (monozygotic and dizygotic twins, non-twin siblings, and unrelated individuals; 93 unique families), all individuals from the same family were randomly assigned to one of two groups of 88 (i.e., split-half cross-validation), with one group being used to train a model that would then be tested on the other (and vice versa). The following approach was then applied to 100 of these random splits of the data to assess the performance of rCPM across different training/testing sets and to build a bagged model that is more robust to overfitting (O’Connor et al., 2020).

1. Calculate leave-one-out (LOO) ISC according to the method described in section 2.4 separately for each group of 88 subjects. Re-calculation of ISC is necessary within each randomly assigned group because calculating an individual’s ISC using data from the 175 subjects across both groups would compromise the independence of the test set by using training subject data to calculate test subjects’ ISC values (Scheinost et al., 2019). We note that ISC estimates have been shown to stabilize at samples of around ∼30 subjects (Pajula & Tohka, 2016), so the magnitudes of the ISC values calculated using either the average of data from 87 or 175 other subjects would not be expected to differ significantly. Next, gISC is calculated for each individual by taking the average of their Fisher *z*-transformed parcel-level ISC values.
2. Identify FC edges whose connectivity strength is most associated with gISC in the 88 subject training group. Spearman partial correlations are performed between each of the 64,620 unique FC edges and gISC, controlling for mean FD in both the concatenated rest and movie day 1 scans, with mean FD calculated as the average of the filtered and uncensored FD traces. Spearman rank correlations are used here and throughout the rest of the paper because the distribution of gISC values exhibited a significant left skew (as did ISC values in general). All edges whose strengths were correlated with gISC at *P* < .01 were retained, although this feature selection step is not strictly necessary for rCPM and exists largely to reduce computational demands (Gao et al., 2019).
3. Fit a regularized linear model using the features (RSFC edges and mean FD measures) identified in the previous step as predictors and gISC as the response. Hyperparameters for this model include alpha (the ridge coefficient), whose optimal value has been shown to be near-zero for true ridge regression, and lambda, which is calculated in an inner fold using the method and code of Gao et al. (2019).
4. Predict gISC for each participant in the test set by multiplying that participant’s RSFC edge strengths for the edges that passed the feature selection step by each edge’s ridge coefficient and adding the intercept. The ridge coefficients, intercept, and optimal lambda value for the present linear model are saved to allow for future bootstrap aggregation (described in the next section).
5. Flip the training and testing groups and repeat steps 1-4.
6. Evaluate model performance by calculating the Spearman partial correlation between the predicted and observed ISC values across participants in the test set, controlling for mean FD from the rest and movie scans.

#### Significance testing the cross-validated models

A permutation scheme was used to assess the statistical significance of the prediction coefficients generated in the 100 split-half iterations. First, the order of the gISC and movie watching mean FD matrices was shuffled such that one participant’s resting state FC matrices and mean FD values were associated with the movie BOLD time courses and mean FD values from a random participant. Steps 1 through 6 from the previous section were then performed and the whole process was repeated 10,000 times. The resulting 10,000 correlation coefficients serve as a null distribution with which the following permutation p-value was calculated (following Finn & Bandettini, 2020): 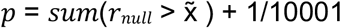, where 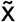 is the median of the 100 correlation coefficients obtained through the true (unshuffled) models.

#### Building and testing the bagged model

The cross-validation paradigm described in the previous two sections was used to ask whether a model trained on data from one group of subjects can predict gISC from RSFC in a novel group of subjects. Importantly, the gISC values used here all come from the same set of stimuli (the Day 1 movie clips). It could then be the case that the CV models rely on stimulus-specific signals and may fail to predict gISC during viewing of a different movie. A bootstrap aggregating, or “bagging,” approach was used to test whether the 200 linear models trained on Day 1 movie watching and resting state data could predict Day 2 gISC (derived from a different set of stimuli) from Day 2 RSFC, as previous work has shown bagged CPM models to be more accurate and more generalizable than their non-bagged counterparts (O’Connor et al., 2020). To construct the bagged model, RSFC edges that passed the *P* < .01 feature selection step in at least 10% (20/200, reflecting the 100 iterations of split-half cross-validation) of iterations were identified, yielding 1,618 edges total. Next, for each of these edges, the ridge coefficients for a given edge across all iterations in which that edge was selected were averaged. The average intercept across the 200 bootstraps was then combined with the average ridge coefficients to yield a singular composite linear model. Each subject’s Day 2 RSFC matrix was then submitted to this model (by multiplying the correlation coefficients of selected RSFC edges by the ridge coefficients and adding the intercept) to generate 176 predicted gISC values, which were then Spearman correlated with the “true” gISC values (calculated through LOO ISC using the full sample), controlling for mean FD in the Day 2 rest and movie scans. To characterize the network-level contribution of edges to the bagged model, the 1,618 ridge coefficients were averaged according to their corresponding edge’s network memberships separately for positive and negative weights.

Code for this rCPM analysis was adapted from github.com/YaleMRRC/CPM (Greene et al., 2020).

### 2.6 Inbound and outbound analyses

To investigate relationships between RSFC and ISC in each of the 360 Glasser parcels, we used two related partial correlation analyses. First, we (Spearman) correlated ISC values for a given parcel with the RSFC coefficients for each edge involving that parcel across participants, controlling for mean FD in the rest and movie scans. This “inbound” analysis describes how ISC in a given parcel is related to that parcel’s resting state functional connections to every other parcel. Next, the “outbound” analysis involves finding the Spearman partial correlation between the RSFC coefficients for each connection between a given parcel and every other parcel and ISC in those other parcels, again controlling for rest and movie scan mean FD. This analysis shows how a given parcel’s resting state functional connections to other parcels are associated with ISC in those parcels. Both the inbound and outbound analyses yield a 360 × 1 map (with one undefined value for the reference parcel) across the brain for every parcel and for each day of data collection, allowing for the visualization of results on the cortical surface.

To identify parcels in which ISC was most positively/negatively associated with RSFC across the brain, the inbound and outbound maps were Fisher *z*-transformed, averaged, and converted back to units of Spearman’s rho. Next, these inbound/outbound averages were grouped by resting state network (RSN) membership according to the Cole-Anticevic Brain-wide Network Partition (Ji et al., 2019) to facilitate visualization of network-level trends.

Because the number of parcels belonging to each Cole/Anticevic RSN is an arbitrary value based on the resolution of the Glasser parcellation, we used bootstrap confidence intervals (CIs) to test whether the averages of the mean inbound and outbound maps within RSN were significantly different from zero. Non-parametric bootstrap CIs were constructed by randomly sampling 176 subjects with replacement, repeating the inbound and outbound analyses, and recalculating the RSN averages 1000 times. To account for multiple comparisons across the twelve RSNs, 95% CIs were Bonferroni corrected using the formula 1 − (alpha/number of RSNs), leading to an effective CI of 99.6% for each RSN average on each day.

The inbound analysis introduced earlier in this section addresses how connectivity from parcel A to parcels B-Z is associated with ISC in parcel A. Although this analysis neatly parallels the dimensionality of the outbound analysis, it ignores how resting state functional connections that don’t involve parcel A (i.e., connectivity *between* parcels B-Z) might be associated with ISC in that parcel. To address this limitation, we used a “full” inbound analysis that correlates all edges of the RSFC matrix with ISC in the reference parcel. We note that this analysis is identical to the initial feature selection step for the rCPM analysis if the gISC values were to be replaced with ISC values for a single parcel.

### 2.7 Spatial permutation tests

Similarity between the Day 1 and Day 2 inbound/outbound maps was quantified using Spearman correlation. Because (1) the number of samples in these correlations is determined by parcellation resolution and (2) spatial autocorrelation is present among adjacent parcels, a spatial permutation test was used to assess the significance of these correlations following the method and code of Váša et al. (2018). Briefly, this test works by inflating one of the spatial maps from each correlation to a sphere, randomly rotating this projection, and calculating the Spearman correlation between the empirical map and the randomly rotated projection. This procedure is repeated until a desired number (here, 10,000) of permutations have been performed, and the final permutation *P* value reflects the number of null permutations for which the resulting Spearman correlation is greater than the observed correlation divided by the total number of permutations. Although recent work has identified that this approach relies upon often unrealistic statistical assumptions (Weinstein et al., 2020), we note that no alternative method is compatible with our effect maps which are undefined at the individual subject level. As such, the present spin method can be seen as the best available significance test for the day-to-day consistency of our observed effects.

### 2.8 Principal component analysis

Principal component analysis (PCA) was performed separately for outbound and (full) inbound matrices from both days of data collection to simplify the matrices into a small number of orthogonal factors. Only the first two PCs of the outbound PCA are visualized in Fig. 4 as these components were sufficient to account for > 50% of the variance in the outbound maps on both days.

**Figure 4.**
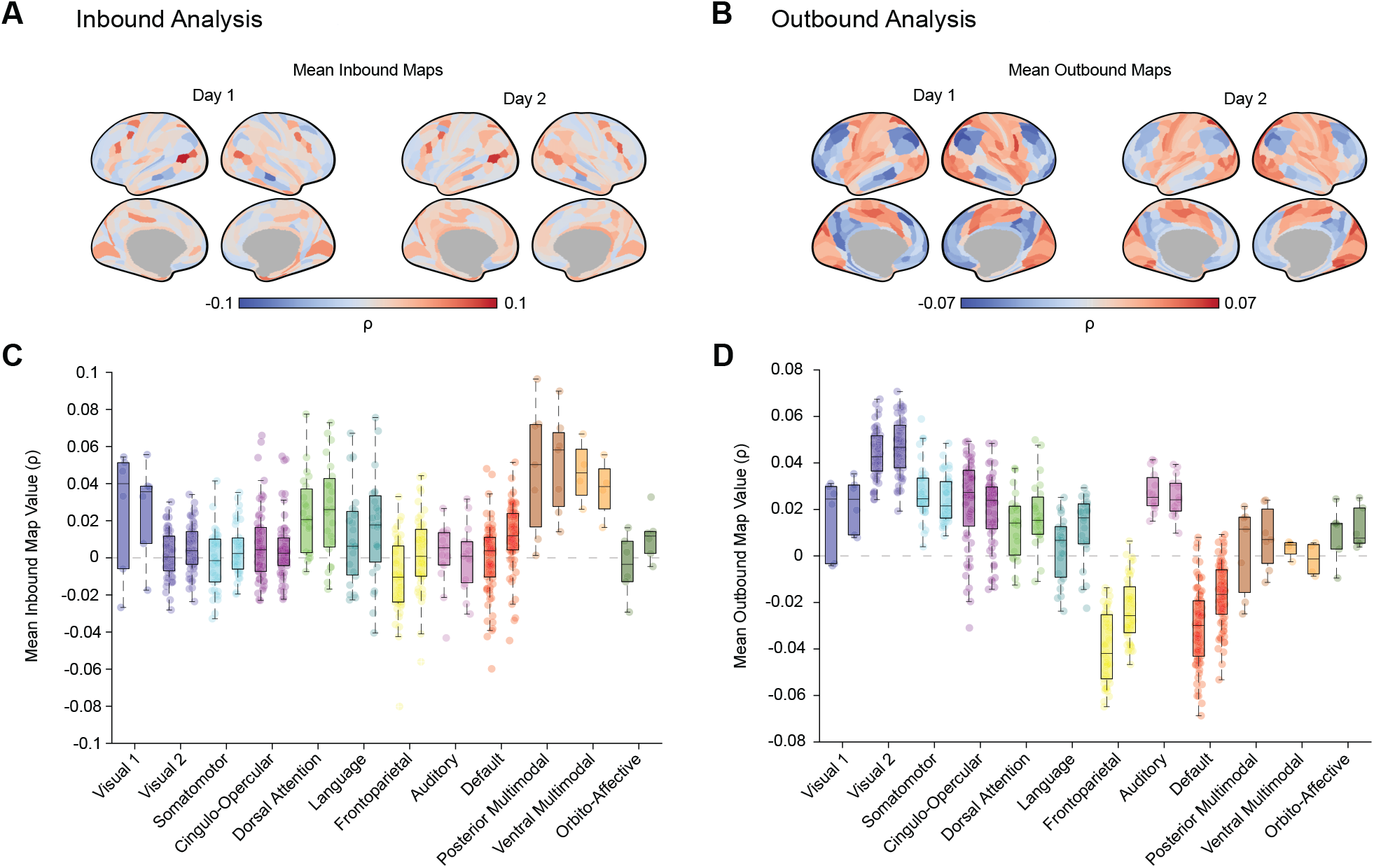
Network-level trends in mean inbound and outbound maps. (A) Each parcel’s values on these surfaces reflect the average of the 359 values comprising its inbound map (e.g., the surfaces in Fig. 3) on each day. (B) Same as (A) for the outbound maps. (C) Boxplots group the parcels from the surfaces in (A) according to their network affiliation, such that the y-coordinates in these plots are the same as the values on the above surfaces. The left and right boxes for each network reflect Day 1 and Day 2 results, respectively. The dorsal attention, posterior multimodal, and ventral multimodal networks had mean inbound values that were significantly greater than zero on both days. Bonferroni-corrected confidence intervals for all other networks contained zero on one or both days. (D) Same as C for the outbound analysis; only visual 2 and frontoparietal networks had average outbound values that were significantly different from zero on both days.

### 2.9 RSFC and ISC topological similarity

One pairwise RSFC similarity matrix was generated for each day of data collection by (Spearman) correlating RSFC profiles (i.e., each subject’s set of 64,620 unique FC edge strengths) from all possible pairs of subjects, leaving two 176 subjects × 176 subjects matrices in which higher values correspond to greater RSFC profile similarity for those two individuals. The same procedure was repeated using each subject’s ISC topologies, again yielding two 176 subjects × 176 subjects matrices. Here, we noted that four subjects, two different subjects on each day, had markedly different ISC topologies from the rest of the sample. These individuals also had the lowest gISC values for their respective movie scans, suggesting minimal engagement with the stimulus. Data from these outlier scans were discarded from the rest of the analyses described in this section for a sample size of 174 subjects, although we note that including these subjects does not change the interpretation of our results. Relatedly, these subjects were not excluded from the previous analyses as they were not identified as outliers in the Fig. 2D and Fig. 3 scatter plots.

**Figure 2.**
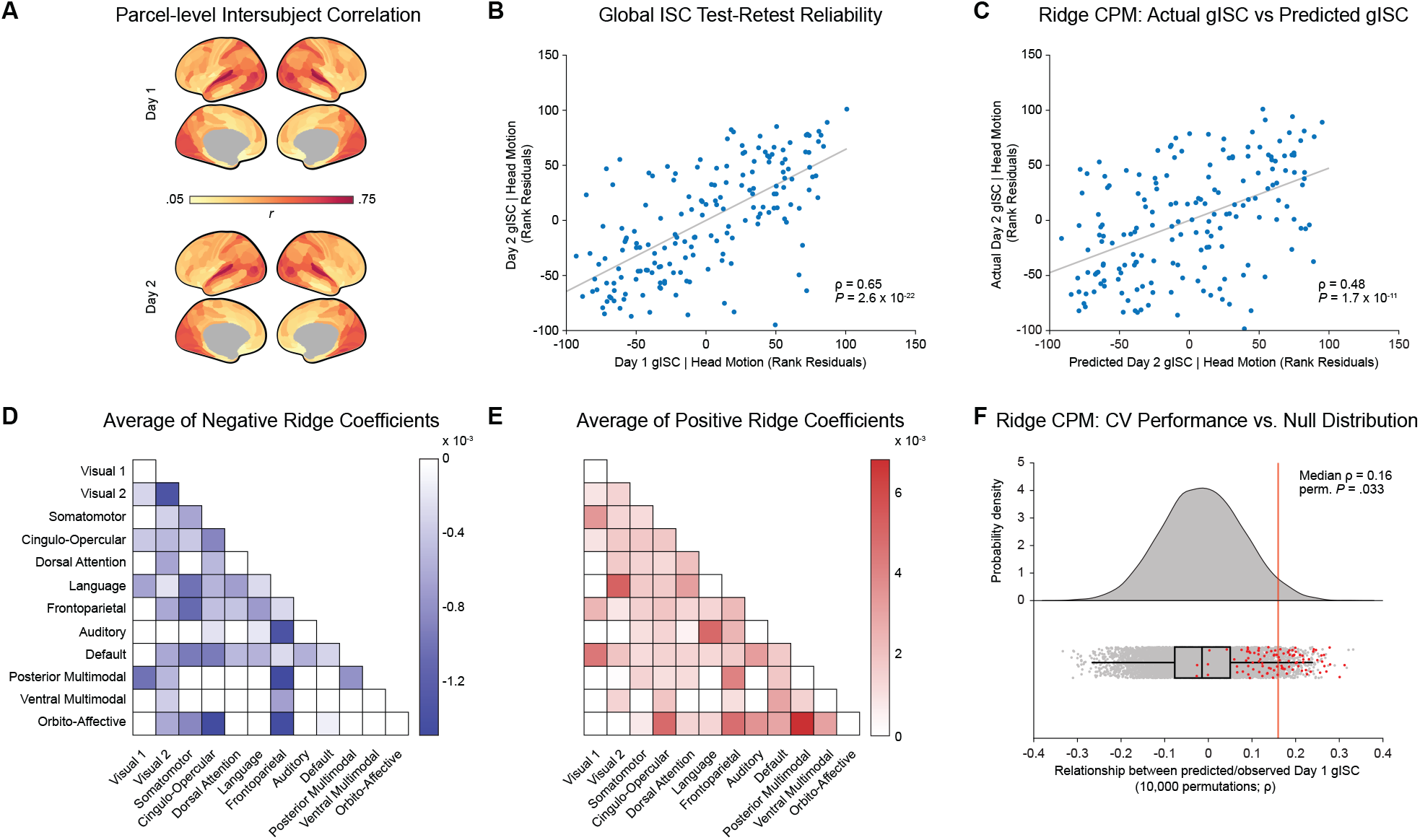
Resting state functional connectivity predicts global intersubject correlation during movie watching in novel stimuli and in novel subjects. (A) Intersubject correlation analysis reveals a non-uniform cortical distribution of shared stimulus-evoked activity that is consistent across Day 1 and Day 2 stimuli (spatial ρ = 0.97, *P*_spin_ < 1 × 10^−4^). (B) Individual differences in gISC were consistent (ρ = 0.65, *P* = 2.6 × 10^−22^) across the Day 1 and Day 2 scans. In this and subsequent scatter plots, each dot represents an individual subject unless noted otherwise. As these plots represent Spearman partial correlations (controlling for head motion), the x- and y-axes are in units of rank residuals so that the slope of the best fit line represents the corresponding Spearman correlation coefficient. (C) A bagged rCPM model trained on Day 1 data predicts Day 2 gISC from RSFC with considerable accuracy (ρ = 0.48, *P* = 1.7 × 10^−11^). (D-E) Heatmaps illustrating that the bagged rCPM model’s negative and positive features were broadly distributed across functional networks. (F) Models trained on Day 1 data from one set of subjects predict Day 1 gISC in held-out subjects at above-chance accuracy. Red dots represent prediction accuracies from the actual 100 CV iterations, whereas gray dots reflect null prediction accuracies from re-running the rCPM analysis using permuted gISC and movie FD values 10,000 times.

**Figure 3.**
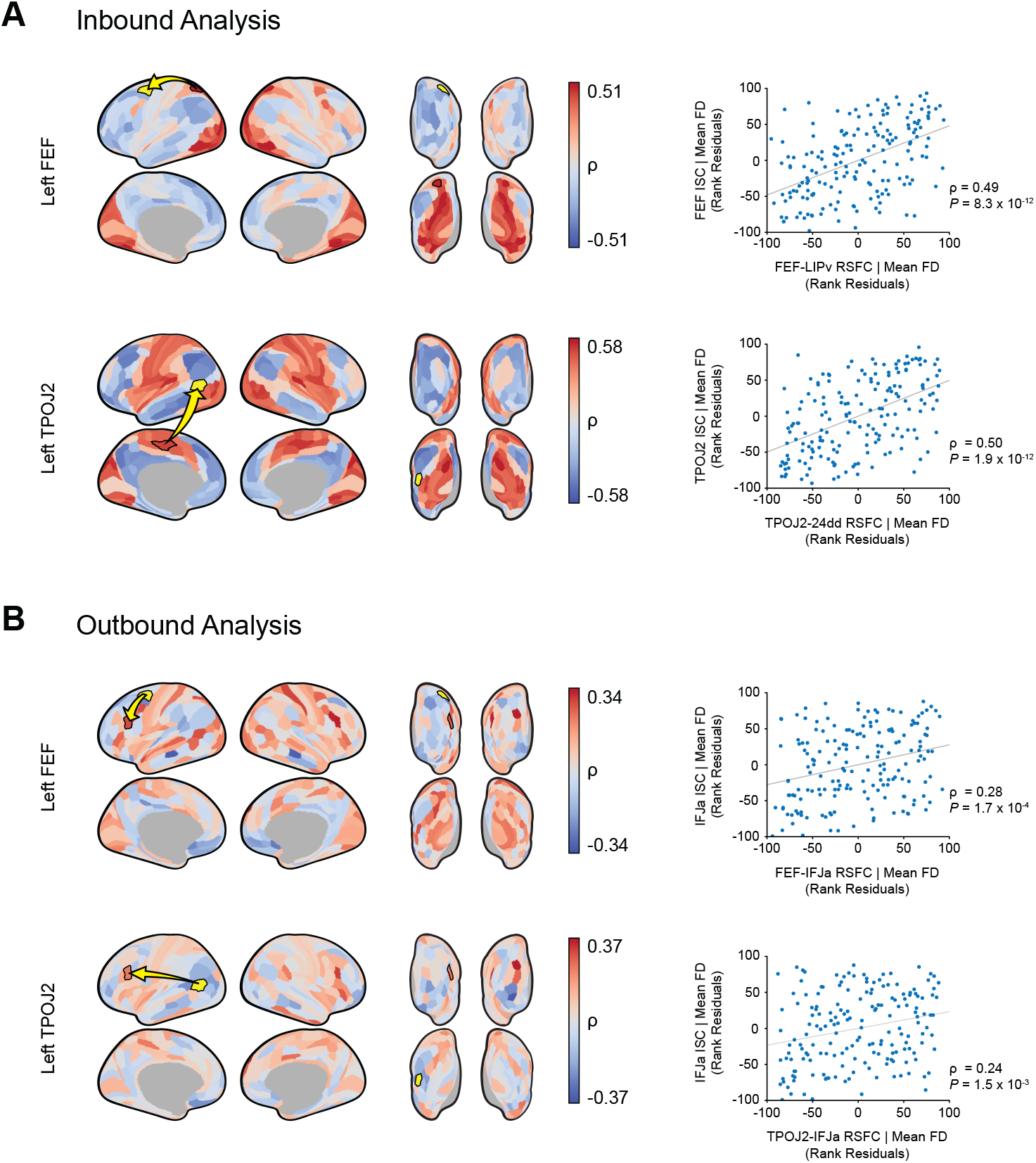
Inbound and outbound relationships between Day 1 RSFC and ISC in FEF and TPOJ2. (A) Inbound analysis: This analysis describes how RSFC between the reference parcel (yellow) and some other parcel (red) is associated with ISC in the <u>reference</u> parcel. Greater resting connectivity from FEF (or TPOJ2) to red-shaded parcels is associated with greater FEF (or TPOJ2) ISC during movie watching. Example inbound maps are shown for parcels covering left FEF (top row) and TPOJ2 (bottom row). Corresponding scatter plots illustrate how the inbound correlation coefficients for the bordered red parcels are calculated. (B) Outbound analysis: This analysis describes how RSFC between the reference parcel (yellow) and some other parcel (red) is associated with ISC in the <u>other</u> parcel. Greater resting connectivity from FEF (or TPOJ2) to red-shaded parcels is associated with greater ISC in those parcels during movie watching. Corresponding scatter plots illustrate how the outbound correlation coefficients for the bordered red parcels were calculated.

Pairwise motion and gISC similarity matrices were also generated by calculating the negative absolute value of the difference between every pair of subjects’ mean framewise displacement for each day’s rest and movie scans as well as their (Fisher *z*-transformed) gISC values. Finally, a Spearman partial correlation was performed between each day’s RSFC and ISC similarity matrices, controlling for pairwise similarity in motion and gISC. A permutation test was used to evaluate the significance of the RSFC-ISC topological similarity relationships. Specifically, the ISC, gISC, and movie watching FD matrices were shuffled together across subjects and ISC/gISC/FD similarities were recalculated. The Spearman partial correlation described above was then performed using the shuffled vectors such that each pair’s mean movie watching FD, gISC, and ISC pattern similarity values were associated with the mean resting state FD and RSFC pattern similarity values from a random pair. This process was repeated 10,000 times to generate the null distributions shown in Fig. 6B.

**Figure 6.**
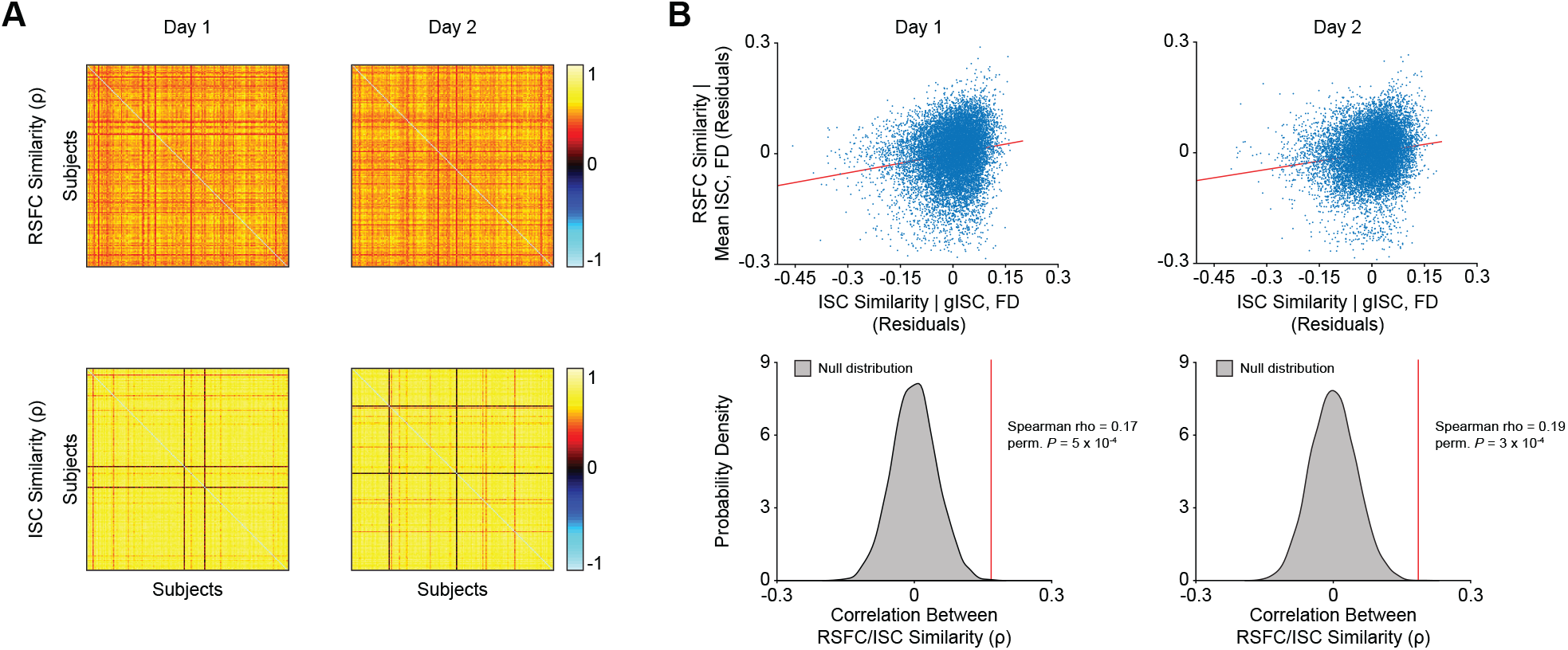
Individuals with more similar resting state connectomes have more similar spatial distributions of ISC. (A) Heatmaps show that RSFC and ISC profiles are generally conserved across individuals. (B) Scatterplots show the positive relationships between RSFC and ISC profile similarity, where each dot reflects one unique pair of subjects. To preserve visual clarity, the x- and y-axes reflect unranked ISC/RSFC similarity values after partialling out the effects of head motion and gISC similarity. The null distributions in the lower row visualize the results of matching one pair’s RSFC similarity with the ISC similarity of a random pair and repeating the Spearman partial correlation 10,000 times. Plotting the strength of the true RSFC/ISC profile similarity correlation coefficients on these distributions reveals that the observed effects are much stronger than would be expected by chance. We note that because unranked values were graphed on the scatterplots, the slopes of the best fit lines only indirectly correspond to the vertical lines on the null distributions.

All analyses were performed in Matlab (R2020a). All cortical surface visualizations were performed with Connectome Workbench (Marcus et al., 2011).

## 3. RESULTS

To predict individual differences in ISC from RSFC, we used publicly available 7T fMRI data collected from 176 Human Connectome Project participants. On each of two days of data collection, all participants completed two movie watching and two resting state runs, each lasting approximately 15 minutes, with one movie run per day featuring clips from major motion pictures and the other consisting of clips from independent short films. All movie clips were viewed only once across the four runs with the exception of a short clip included at the end of every run for test-retest purposes. Functional data were preprocessed, averaged into 360 cortical regions of interest according to the Glasser parcellation (Glasser et al., 2016), and concatenated by scan type and date to yield one composite resting state and one composite movie watching scan for each day of data collection. A 360 parcels × 360 parcels × 176 subjects × 2 days RSFC matrix was then generated by (Pearson) correlating resting state activity time courses from all pairs of cortical parcels (Fig. 1A, step 1) for each subject and for each day. Next, for each of the two composite movie scans, each participant’s activity time course in a given parcel was (Pearson) correlated with that parcel’s average BOLD signal time course across all 175 other participants. This process was repeated across all parcels to create a 176 subjects × 360 parcels × 2 days ISC matrix in which higher values correspond to more typical movie-evoked brain activity.

**Figure 1.**
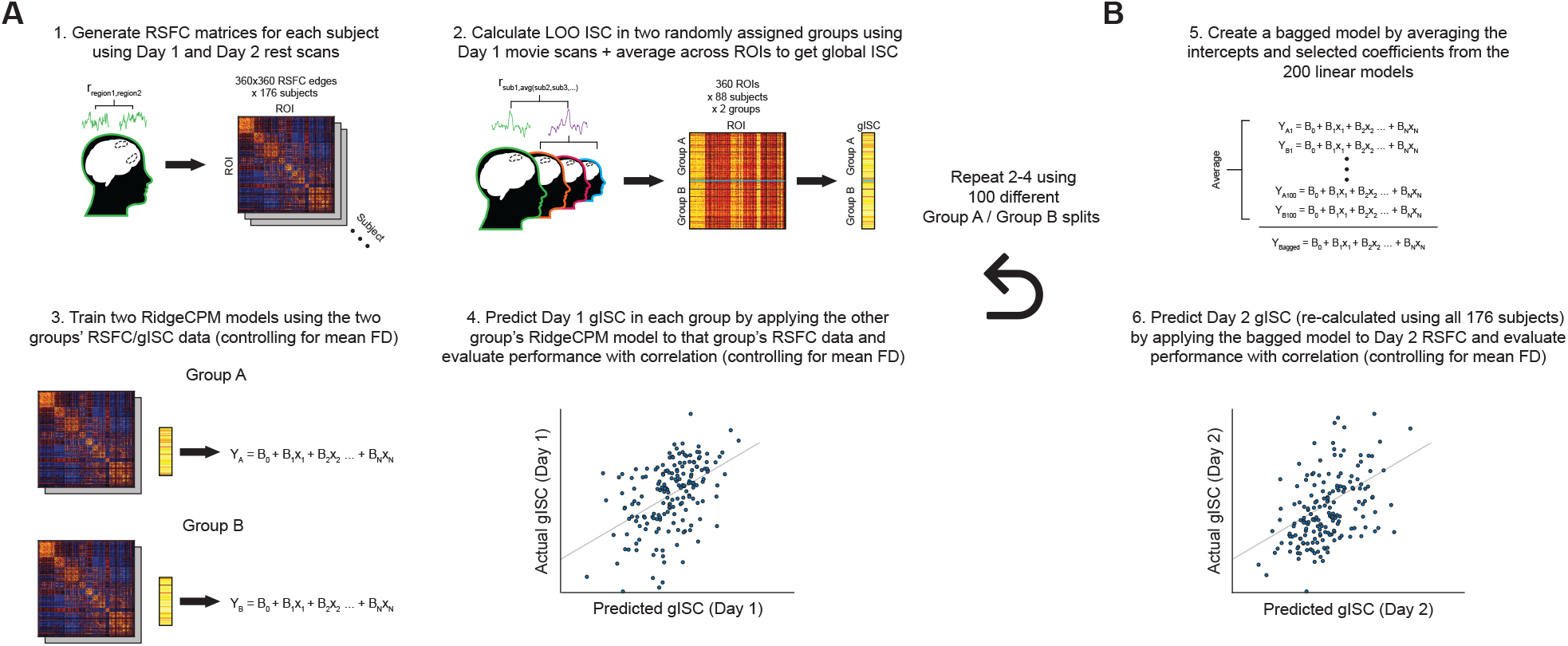
Ridge CPM pipeline. (A) Predicting gISC from RSFC in held-out subjects: RSFC matrices were created by (Pearson) correlating activity time courses from all pairs of parcels for each individual and day of data collection. Subjects were divided into two commensurate groups, and ISC values for every parcel were calculated for each subject as the Pearson correlations between their activity time course in a given parcel and that of the group average. These 360 parcel-wise values were averaged to yield 1 gISC value per subject per day of data collection. A model was then trained to predict gISC values from RSFC data in one group and tested on data from the held-out group, with model performance being evaluated as the Spearman partial correlation between predicted and actual Day 1 gISC values, controlling for head motion. This procedure was repeated 99 more times using different train/test group splits, yielding 200 linear models and 100 correlation coefficients reflecting prediction accuracies. (B) Predicting gISC from RSFC in held-out stimuli: The 200 linear models were aggregated into one composite model which was tested on Day 2 gISC and RSFC data from all 176 subjects. Once again, model accuracy was evaluated using the Spearman partial correlation between predicted and actual Day 2 gISC values, controlling for Day 2 head motion.

### 3.1 Functional connectivity at rest predicts a global measure of normative movie-evoked activity

Ridge connectome-based predictive modeling (rCPM) was used to predict each participant’s average ISC across all parcels. This cortex-wide measure, which we refer to as global ISC (gISC), serves as a low-dimensional marker of brain synchronization, or how similar an individual’s temporal patterns of neural activity during movie watching are to the rest of the group’s. Our use of this measure is motivated by its simplicity as well as by previous work showing global relationships between ISC and behavior (Gruskin et al., 2020). RCPM is a prediction method that works by identifying RSFC edges (or “features”) whose connectivity strength is associated with a phenotype of interest (here, gISC) in a training set and using those features to fit a regularized linear model that can be tested on data from held-out participants (Gao et al., 2019). We used rCPM to ask two related questions: to what extent can a model trained on data from one set of subjects watching one set of movie clips predict gISC in (1) a different set of subjects watching the same clips, and (2) the same subjects watching a different set of clips.

Using only the Day 1 rest and movie data, we performed 100 iterations of rCPM with split-half cross-validation (CV) to predict gISC from RSFC in unseen subjects. To keep training and testing data independent, gISC was calculated separately for each group (Fig. 1A, step 2). Each rCPM iteration yielded two linear models (one for each split-half; Fig. 1A, step 3) and one correlation coefficient reflecting the Spearman partial correlation between predicted gISC and observed gISC (controlling for average motion in the rest and movie scans; Fig. 1A, step 4). Spearman correlations were used here and throughout the paper due to a significant left skew in the distribution of gISC. The linear models from all 100 iterations were then combined into a single model following a bootstrap aggregating (“bagging”) approach that has been shown to reduce overfitting and improve model accuracy (Fig. 1B, step 5; O’Connor et al., 2020). This bagged model was then used to predict Day 2 gISC from Day 2 RSFC (Fig. 1B, step 6).

Parcel-level ISC (averaged across individuals; Fig. 2A) and individual-level gISC (averaged across parcels; Fig. 2B) were consistent across the two days, as reflected by Spearman correlations of ρ = 0.97 (*P*_spin_ < 1 × 10^−4^) and ρ = 0.65 (*P* = 2.6 × 10^−22^) respectively. We note that the ρ = 0.65 test-retest reliability value serves as a sort of upper bound on prediction performance as it represents how well gISC can predict itself across the different stimuli. Because the reliability of gISC was calculated while controlling for head motion (quantified by mean framewise displacement; mean FD) from both movie scans, rank residuals were used to create the scatterplots shown in Fig. 2 and elsewhere in the paper when appropriate (although plots showing raw values can be found in Supplementary Fig. 1).

The bagged model was able to predict Day 2 gISC from Day 2 RSFC with significant accuracy (ρ = 0.48, *P* = 1.7 × 10^−11^; Fig. 2C) indicating that an individual’s ability to exhibit normative stimulus-evoked activity is closely related to their brain’s intrinsic functional architecture. To visualize the functional distribution of RSFC edges that contributed to the model’s predictive performance, we averaged positive and negative edges separately according to resting state network (RSN) membership as defined by the Cole-Anticevic Brain-wide Network Partition (Ji et al., 2019). Predictive edges were widely distributed across RSNs according to no obvious pattern, although positive relationships between RSFC and gISC were more pronounced than negative relationships (as seen in the scales of Figs. 2D-E).

To examine how models trained on data from one group of subjects can predict gISC in held-out individuals watching the same stimulus, we plotted the prediction accuracies obtained from the 100 CV iterations (Fig. 2F, red dots) against a distribution of null accuracies (Fig. 2F, gray dots) generated by permuting gISC values across subjects and re-running the rCPM pipeline 10,000 times. As shown in Fig. 2F, the median CV accuracy was significantly greater than would be expected by chance (median ρ = 0.16, permutation *P* = .033).

### 3.2 Inbound and outbound analyses characterize region-specific RSFC-ISC relationships

In the previous section, we demonstrated that an individual’s RSFC profile is predictive of their cortex-wide ISC during movie watching. Next, to investigate region-specific RSFC-ISC relationships, we introduce the related “inbound” and “outbound” analyses. The inbound analysis, demonstrated in Fig. 3A using Day 1 data, describes how ISC in a given parcel is associated with that parcel’s resting connectivity to every other parcel (raw values are shown in Supplementary Fig. 2). The intuition behind and results of this analysis are first exemplified using the left frontal eye field (FEF), a parcel chosen for its circumscribed role in the well-studied saccade pathway (Felleman & Van Essen, 1991; Paus, 1996). To create the brain maps shown in the top panel of Fig. 3A, left FEF ISC was correlated with RSFC between left FEF and the other 359 cortical parcels while controlling for head motion from the corresponding rest and movie scans. The resulting correlation coefficients were then projected onto an inflated cortical surface for visualization, revealing that individuals whose left FEF was more connected to visual and parietal areas at rest exhibited more typical FEF activity during movie watching. We note that this finding is consistent with the FEF’s known inputs from visual cortex and parietal attention areas (Felleman & Van Essen, 1991; Paus, 1996).

Having demonstrated that the inbound analysis reveals expected RSFC-ISC relationships in a relatively unimodal and well-studied area, we next sought to apply the same analysis to a parcel in the temporo-parieto-occipital junction (TPOJ), a multimodal area in the temporoparietal junction/posterior superior temporal sulcus region whose functioning is less understood but has been implicated in sensory integration during movie watching (Lerner et al., 2011; Patel et al., 2019). In this parcel, greater ISC was associated with increased connectivity to unimodal sensory (visual, auditory, somatomotor) cortices and decreased connectivity to higher order areas (e.g., angular gyrus, dorsolateral prefrontal cortex; dlPFC) at rest (Fig. 3A). Although only Day 1 results are visualized in Fig. 3, Spearman spatial correlations confirmed that the inbound maps for both left FEF and TPOJ2 were consistent across both days of data collection (spatial ρ = 0.91, *P*_*spin*_ < 1 × 10^−4^ and spatial ρ = 0.98, *P*_*spin*_ < 1 × 10^−4^ respectively).

While the inbound analysis describes how a brain region’s functioning (ISC) is related to its intrinsic connectivity (RSFC) to other regions, a distinct but similarly informative relationship is how a region’s RSFC to other areas is associated with ISC in those areas. To quantify this “outbound” relationship, we calculated the Spearman partial correlation between the RSFC edge strengths for all connections involving a given parcel and ISC in every other parcel. The FEF outbound map therefore illustrates that greater RSFC between FEF and dlPFC is associated with greater ISC in dlPFC (Fig. 3B), again replicating the known hierarchical relationships between these two areas (Arkin et al., 2020; Corbetta et al., 2008). The FEF and TPOJ2 outbound maps were also found to be consistent across the different days of data collection (spatial ρ = 0.84, *P*_*spin*_ < 1 × 10^−4^ and spatial ρ = 0.87, *P*_*spin*_ < 1 × 10^−4^ respectively).

Classical theories of sensory information processing place cortical areas on a spectrum ranging from specialized to integrative (Mesulam, 1998). To identify integrative hubs whose normal functioning during movie watching is most associated with greater intrinsic connectivity to the rest of the brain, we visualized the average of each parcel’s inbound map and grouped these average values by RSN affiliation to illustrate network-level trends in Fig. 4. Across both days, parcels comprising the TPOJ exhibited the highest average inbound values, followed by several dlPFC parcels, indicating that greater ISC in these regions was most associated with greater RSFC to the rest of the brain (Fig. 4A).

At the network level, the posterior and ventral multimodal (PMM/VMM) networks displayed the highest average inbound values (Fig. 4C). In addition to the dorsal attention network (DAN), these were also the only RSNs whose average values were found to be significantly different from zero on both Day 1 (PMM mean ρ = 0.048, 99.6% bootstrap CI [0.02, 0.077]; VMM mean ρ = 0.046, 99.6% bootstrap CI [0.011, 0.082]; DAN mean ρ = 0.023, 99.6% bootstrap CI [0.001, 0.047]) and Day 2 (PMM mean ρ = 0.051, 99.6% bootstrap CI [0.041, 0.083]; VMM mean ρ = 0.037, 99.6% bootstrap CI [0.01, 0.071]; DAN mean ρ = 0.025, 99.6% bootstrap CI [0.023, 0.055]). Like the integrative parcels described above, this result suggests that parcels comprising these multimodal networks encode more normative stimulus-evoked information during movie-watching when they are more globally connected to other brain regions at rest.

Complementary analyses were next performed using the outbound maps to identify source-like parcels to which greater connectivity at rest was associated with greater ISC during movie watching (Figs. 4B and 4D). The visual 2 and auditory networks exhibited the highest and second-highest average outbound values on both Day 1 (mean visual 2 ρ = 0.044, 99.6% bootstrap CI [0.017, 0.068]; mean auditory ρ = 0.027, 99.6% bootstrap CI [-0.001, 0.055]) and Day 2 (mean visual 2 ρ = 0.047, 99.6% bootstrap CI [0.021, 0.06], mean auditory ρ = 0.025, 99.6% bootstrap CI [0.026, 0.075]). On the other hand, the frontoparietal and default mode networks (FP/DMN) exhibited negative average outbound relationships on Day 1 (FPN mean ρ = −0.04, 99.6% bootstrap CI [-0.054, −0.022]; DMN mean ρ = −0.031, 99.6% bootstrap CI [-0.049, −0.009]) and Day 2 (FPN mean ρ = −0.023, 99.6% bootstrap CI [-0.055, −0.023]; DMN mean ρ = −0.017, 99.6% bootstrap CI [-0.022, 0.018]). Although the Day 1 auditory and Day 2 DMN CIs contained zero, this overall pattern of results indicates that for the average parcel, greater connectivity at rest to unimodal sensory areas (i.e., auditory and visual networks) was most associated with greater ISC during movie watching, while the opposite was true regarding greater connectivity to higher-order networks (i.e., FPN and DMN).

### 3.3 Principal component analysis reveals low-dimensional embedding of inbound and outbound maps

The average outbound maps shown in the previous section suggest that the more connected a parcel is to visual and auditory regions at rest, the more normative stimulus-evoked activity it exhibits during movie watching. However, this interpretation is complicated by the fact that these sensory regions are among the most synchronized during movie watching, making it likely that they have stronger RSFC-ISC relationships which would contribute more to the average outbound maps. It could then be the case that greater ISC in a given parcel is associated more with greater connectivity to functionally similar parcels (e.g., visual to visual, default mode to default mode) than to sensory parcels in general (e.g., any parcel to visual). A similar ambiguity arises from the average inbound maps, as ISC in parcels with low average inbound values could be correlated in either strong and localized or weak and distributed patterns with RSFC. To explore the underlying structure of the 360 inbound and outbound maps and resolve these ambiguities, we used principal component analysis (PCA), a dimension reduction technique that preserves more of the information in the original feature space than the averages shown in section 3.2.

PCA of the outbound maps revealed that two PCs (Fig. 5A, lower row) were sufficient to account for a majority of the outbound map variance on both days, describing ∼30% and ∼20% of variance, respectively (Fig. 5B). Here, parcels with higher (redder) scores in the upper rows have outbound maps that look like the PCs shown in the lower rows. Relatedly, given the symmetrical nature of the outbound and inbound matrices, parcels with stronger loadings on these PCs have inbound maps that look more like the cortical maps shown in the upper rows.

**Figure 5.**
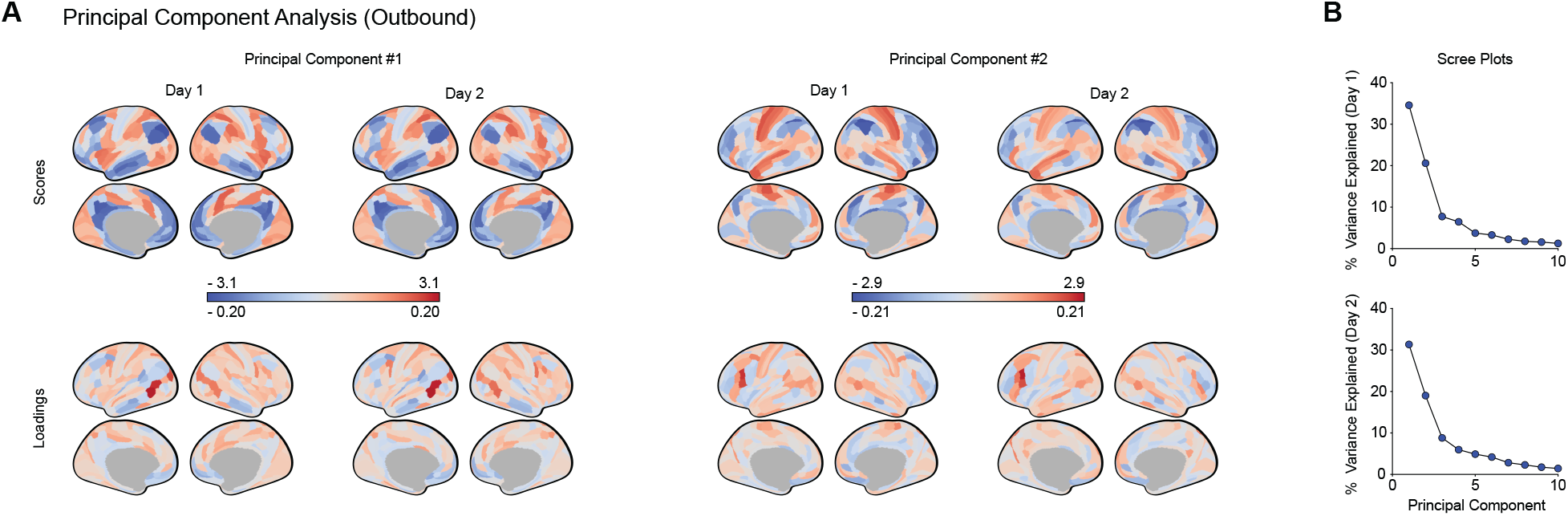
Principal component analysis of the outbound maps. (A) Cortical surfaces display the scores and loadings for the first two principal components (PCs) of the outbound maps. Parcels in the lower row of surfaces that appear redder have inbound maps that look more like the surfaces in the upper row. Due to the symmetry of the inbound/outbound maps, redder parcels in the upper row of surfaces have outbound maps that look more like the surfaces in the lower row. (B) Scree plots show that the majority of outbound map variance can be accounted for by two PCs, with the percentage of variance explained tapering off after the third PC on both days.

The first PC was most positively expressed in visual, cingulo-opercular, and dorsal attention parcels, and was negatively expressed in DMN parcels. This PC loaded most strongly onto left TPOJ, but default mode parcels loaded either negatively or only weakly positively on this component. In other words, default mode parcels exhibited greater ISC when they were more connected to other default mode (and less connected to visual) parcels in this PC. This result suggests that greater ISC is indeed associated with greater RSFC to functionally similar, not just sensory, regions.

The second PC, most positively represented in the superior temporal sulci (STS) and somatomotor cortices and negatively represented in FPN parcels, loaded most heavily onto left dlPFC. Although TPOJ and dlPFC had among the highest average inbound values in Fig. 4, the former’s loading onto PC1 and the latter’s loading onto PC2 suggests that greater ISC in these two integrative areas is most associated with connectivity to separate sets of regions. This difference is especially apparent when considering connectivity to STS, which was positively associated with dlPFC ISC (in PC2) but negatively associated with ISC in TPOJ (in PC1).

While the inbound analysis introduced in section 3.2 neatly parallels the dimensionality of the outbound analysis, it only captures how ISC in parcel A is associated with its RSFC *to* parcels B-Z, ignoring RSFC *between* parcels B-Z. To fill this gap, results from a version of the inbound PCA that relates RSFC across all edges with ISC are visualized in Supplementary Fig. 3, which shows that ISC in different functional systems is most associated with notably different full RSFC patterns.

### 3.4 Subjects with more similar RSFC fingerprints share more similar spatial patterns of ISC

Our analyses thus far have focused on relationships between the magnitudes of RSFC and ISC across participants. While most studies on individual differences in ISC have similarly concentrated on ISC magnitude, the spatial pattern of ISC across a person’s brain likely reflects additional individual-specific aspects of brain function. For example, two subjects with identical FEF lesions might have different *absolute* levels of global and FEF ISC due to differences in arousal during movie watching, but both subjects would be expected to exhibit the same *relative* ISC deficit in FEF compared to their other brain regions. Such an emphasis on topological patterns is apparent in functional connectome fingerprinting studies, which have shown that individuals can be identified by their unique set of (relative) RSFC edge strengths (Finn et al., 2015). However, the relevance of these whole-brain RSFC profiles to spatial patterns of ISC is still unclear.

To investigate whether RSFC profiles are related to how ISC, independent of its overall magnitude, is distributed across the brain, we used a pairwise similarity analysis to determine if individuals with more similar functional connectomes also shared more similar spatial patterns of ISC. First, we correlated RSFC and ISC profiles separately across participants to generate the similarity matrices shown in Fig. 6A. As expected, patterns of RSFC and ISC were largely conserved across participants (Day 1/Day 2 RSFC mean pairwise similarity ρ = 0.52/0.53, std. = 0.015/0.012; ISC mean pairwise similarity ρ = 0.74/0.74, std. = 0.017/0.021). On each day, the ISC topologies for two individuals were markedly different compared to the rest of the sample (seen as dark stripes in the ISC similarity heatmaps in 6A, different individuals on each day), so these four outlier movie scans were discarded from the following analysis. Next, we correlated the pairwise ISC and RSFC similarity values, controlling for similarity in gISC and rest and movie scan head motion. Across both days, we found that participant pairs with more similar RSFC profiles also shared more similar spatial patterns of ISC (Day 1 ρ = 0.17, permutation *P* = 5 × 10^−4^; Day 2 ρ = 0.19, permutation *P* = 3 × 10^−4^; Fig. 6B, upper row; raw values shown in Supplementary Fig. 4), regardless of their similarities in ISC magnitude and head motion. Significance testing for this effect was performed by matching the RSFC similarity of one pair with the ISC and gISC similarities of a random pair, re-running the previous analysis 10,000 times, and calculating a permutation *P* value using the true and null correlation coefficients. The null distributions generated for this test are shown in Fig. 6B (lower row) and illustrate that the topological similarity relationships are significantly stronger than would be expected by chance.

## 4. DISCUSSION

Here, we extend research into the neural processing of naturalistic stimuli by showing that an individual’s brain connectivity at rest is associated with their degree and distribution of normative movie-evoked brain activity. First, we demonstrated that distributed networks of RSFC edges can be used to predict global ISC across subjects and scanning sessions. Next, we explored regional RSFC-ISC relationships, finding that greater ISC in multimodal parcels is related to higher RSFC to unimodal sensory cortex and lower RSFC to frontoparietal and default mode networks. Finally, we showed that individuals with more similar RSFC fingerprints share more similar spatial patterns of ISC. Taken together, these results reveal new relationships between intrinsic brain connectivity and task-driven function while opening up additional lines of inquiry for the study of idiosyncratic responses to naturalistic stimuli.

Individual differences in ISC magnitude have previously been associated with behavioral traits such as top-down attentional control (Campbell et al., 2015). Building upon this literature, we used rCPM to identify a network of RSFC edges (i.e., the 1,618 RSFC edges included in the bagged model) whose combined strengths accounted for a sizable proportion of between-subject variance in a global measure of ISC. Extending the work of Campbell and colleagues, this RSFC network could reflect a general signature of attentional ability, similar to networks identified by Rosenberg et al. (2016, 2020). In this case, individuals who express this RSFC profile more so than others would be expected to attend more to movie clips, leading to increased levels of ISC. At the parcel level, we would expect ISC in different parcels to be associated with the same 1,618 RSFC edges that comprise the predictive network, as greater attention to the movie clips should lead to distributed increases in ISC (Ki et al., 2016; Regev et al., 2019; Song et al., 2020). Instead, we observed that ISC in different parcels was associated with many different patterns of RSFC. Specifically, 10+ principal components were needed to account for a majority of the full inbound matrix variance shown in Fig. S3. Given this result, the RSFC network might instead represent a functional architecture that is more conducive to bottom-up information transfer, independent of top-down influences. In actuality, this network likely represents a combination of top-down and bottom-up effects, and future studies that simultaneously evaluate relationships between RSFC, ISC, and behavioral traits like attentional control will provide valuable insight into the cognitive implications of the present findings.

In addition to identifying RSFC edges that predict global ISC, we also characterized region-specific RSFC-ISC relationships through our inbound and outbound analyses. First, our inbound analysis showed that the resting state connections most associated with ISC in a given parcel are not simply a function of its network affiliation. For example, although the FEF parcel studied here was shown to be more connected to cingulo-opercular areas than to visual cortex at rest (Ji et al. 2019), our inbound analysis showed that typical activity in this parcel during movie-watching was more strongly associated with its connectivity to visual rather than cingulo-opercular cortex. Furthermore, although RSFC derived from bivariate correlation is inherently undirected, comparing the inbound and outbound maps for a given parcel allows for the generation of directional hypotheses. That visual-FEF connectivity is more associated with FEF ISC than visual cortex ISC suggests that, consistent with established models of the visual processing hierarchy (Felleman & Van Essen, 1991), stimulus-related information is more likely to flow from visual cortex to FEF. This expected finding, as well as the stability of these results across different rest and movie scans, motivates the use of inbound/outbound analyses to investigate how connectivity relates to function in more arcane regions and functional systems, as we demonstrated with the TPOJ.

After showing how the inbound and outbound analyses reveal stable RSFC-ISC relationships in specific regions, we next investigated network-level trends in these relationships. By averaging across the inbound maps, we found that parcels in the ventral and posterior multimodal networks (located in temporoparietal cortex) exhibited the highest average inbound map values. In other words, these parcels were more synchronized when they were more positively connected to the rest of the brain. This finding is consistent with previous work showing that these posterior multimodal regions consolidate audiovisual information across long temporal windows (Baldassano et al., 2017; Ji et al., 2019; Lerner et al., 2011) and suggests that the integrative function of these regions is related to their distributed intrinsic connectivity. Complementing this result, we found that parcels in visual and auditory networks exhibited some of the greatest average outbound values, indicating that greater connectivity to these areas at rest was associated with greater ISC. This likely reflects the fact that auditory and visual areas are among the most synchronized during movie-watching and can be thought of as sources of stimulus-driven signal. Alternatively, connectivity to frontoparietal and default mode parcels, which tend to be among the least synchronized by movies, was on average inversely related to ISC. Subsequent PCA of the outbound maps clarified this result by showing that connectivity to default mode parcels was negatively associated with TPOJ ISC but positively associated with ISC in default mode parcels themselves. That these parcels exhibited more typical activity when they were more connected to functionally similar (and less connected to dissimilar) areas at rest is consistent with the idea that more modular cortical organization is associated with more efficient information flow (Wig, 2017). Lastly, the loading of left dlPFC and TPOJ onto different PCs indicates that ISC in both areas is closely related to their RSFC to different sets of parcels, suggesting that these areas may play important but distinct roles in the processing of naturalistic stimuli.

Beyond informing theories of cortical information processing, our results have implications for the understanding of circumscribed ISC deficits reported in patient populations. Studies that link symptom severity or clinical diagnoses with lower ISC in a particular brain region have tended to interpret these relationships as reflecting an intra-regional impairment. Our inbound/outbound findings provide evidence for the alternative hypothesis that low ISC in a particular area may be driven by pathological alterations in inter-regional connectivity rather than dysfunction in the region itself. Future work outlining the causal direction of these RSFC-ISC relationships will inform the clinical applications of naturalistic fMRI (Eickhoff et al., 2020), as whether atypical activity in a region is self-limited or a consequence of its connectivity to other areas is an important consideration in developing targeted therapies (Fox et al., 2012).

Although naturalistic imaging studies have tended to focus on individual differences in the magnitude of ISC, it is important to note that this measure is susceptible to session effects that may obscure meaningful trait-related variance. For example, an attentive subject scanned at the end of a long day may exhibit less ISC than a more distractible individual scanned when they are most alert. The low test-retest reliability of activation magnitudes is a general issue in task fMRI research (Elliott et al., 2020). However, by asking “where” task activations occur instead of “how much” activation is present, researchers have been able to more reliably elicit individual differences and excise state-dependent noise from their analyses (Kragel et al., 2020). Moreover, individuals’ activation maps derived from modeled responses to difficult tasks have been accurately predicted from their RSFC profiles, suggesting that unique spatial activation patterns are intimately related to variation in the brain’s intrinsic wiring (Cole et al., 2016; Tavor et al., 2016). Extending this line of work to model-free naturalistic paradigms, we found that subjects with more similar resting state connectomes shared more similar spatial patterns of ISC during movie watching, independent of their similarity in ISC magnitude. This finding indicates that spatial distributions of ISC capture subject-specific information that ISC magnitudes alone do not, echoing recent reports that have established multivariate patterns as superior detectors of individual differences (Kragel et al., 2020).

Several limitations of this study constrain the interpretation and generalizability of our results. First, the short (3-5 minute) clips used here may have failed to adequately engage cortical areas with longer temporal receptive windows (Baldassano et al., 2017). Future studies that utilize continuous stimuli may be better able to characterize RSFC-ISC relationships in these areas. Second, functional brain topologies have been shown to differ meaningfully across individuals and task states (Gratton et al., 2018; Salehi et al., 2020). Subjects identified as having lower RSFC and/or ISC in our analyses may instead have functional topologies that are less represented by the Glasser parcellation, complicating the interpretation of our inbound/outbound results. Finally, our use of Spearman correlations to link RSFC and ISC presents its own set of limitations. Some RSFC-ISC relationships may be non-monotonic, such that both hypo- and hyper-connectivity between a pair of regions is associated with lower ISC. The correlation method used here also prevents us from assessing the unique contribution of single RSFC edges to parcel-level ISC, as would be possible with multiple regression.

Despite these limitations, this study constitutes an important step towards characterizing the complex relationships between normative stimulus-evoked activity and the brain’s intrinsic functional architecture. By linking resting state connectivity with task-driven function at the parcel, network, and cortex-wide levels, these findings further develop our understanding of sensory information processing and anchor individual differences in ISC in the resting state connectome. Additional research into these connectivity-activity relationships will be instrumental in exploring the cognitive and perceptual relevance of resting state fMRI and contextualizing new discoveries from the growing field of naturalistic imaging.

## DATA AND CODE AVAILABILITY

The raw HCP data used for this project can be downloaded from ConnectomeDB (db.humanconnectome.org). Code for all analyses can be found in the following Github repository: https://github.com/davidgruskin/hcp_rsfc_isc.

## CRediT STATEMENT

David C. Gruskin: Conceptualization, Methodology, Formal Analysis, Visualization, Writing - Original Draft, Writing - Review & Editing Gaurav H. Patel: Conceptualization, Writing - Review & Editing, Supervision

## CITATION DIVERSITY STATEMENT

Recent work in several fields of science has identified a bias in citation practices such that papers from women and other minority scholars are under-cited relative to the number of such papers in the field (Caplar et al., 2017; Dion et al., 2018; Dworkin et al., 2020; Maliniak et al., 2013; Mitchell et al., 2013). Here we sought to proactively consider choosing references that reflect the diversity of the field in thought, form of contribution, gender, race, ethnicity, and other factors. First, we obtained the predicted gender of the first and last author of each reference by using databases that store the probability of a first name being carried by a woman (Dworkin et al., 2020; Zhou et al., 2020). By this measure (and excluding self-citations to the first and last authors of our current paper), our references contain 5.88% woman(first)/woman(last), 3.92% man/woman, 29.41% woman/man, and 60.78% man/man. This method is limited in that a) names, pronouns, and social media profiles used to construct the databases may not, in every case, be indicative of gender identity and b) it cannot account for intersex, non-binary, or transgender people. Second, we obtained predicted racial/ethnic categories of the first and last author of each reference by databases that store the probability of a first and last name being carried by an author of color (Ambekar et al., 2009; Sood & Laohaprapanon, 2018). By this measure (and excluding self-citations), our references contain 6.16% author of color (first)/author of color(last), 15.95% white author/author of color, 24.05% author of color/white author, and 53.84% white author/white author. This method is limited in that a) names and Florida Voter Data to make the predictions may not be indicative of racial/ethnic identity, and b) it cannot account for Indigenous and mixed-race authors, or those who may face differential biases due to the ambiguous racialization or ethnicization of their names. We look forward to future work that could help us to better understand how to support equitable practices in science.

## ACKNOWLEDGEMENTS

We thank Monica Rosenberg and Ajay Nadig for feedback on earlier versions of this project. Author D.C.G. was supported by a NIH MSTP training grant (T32GM007367). Author G.H.P. was supported by the NIMH (K23MH108711, R01MH121790, and R01MH123639) and by a David Mahoney Neuroimaging Grant from the DANA Foundation. Data were provided by the Human Connectome Project, WU-Minn Consortium (Principal Investigators: David Van Essen and Kamil Ugurbil; 1U54MH091657) funded by the 16 NIH Institutes and Centers that support the NIH Blueprint for Neuroscience Research; and by the McDonnell Center for Systems Neuroscience at Washington University. The authors have no competing interests to declare.

---

**Figure S1.**
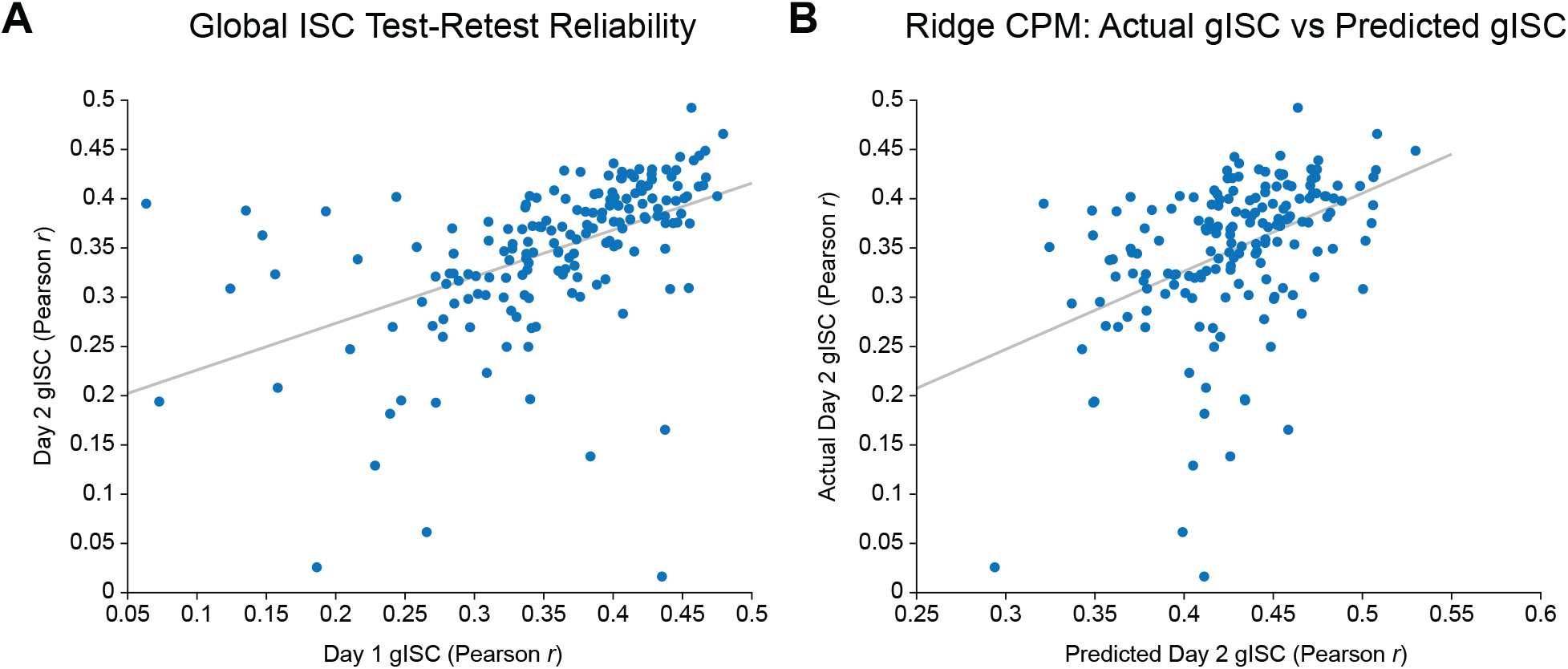
Raw gISC scatterplots. (A) Individual differences in gISC were consistent across the Day 1 and Day 2 scans, but low gISC values on one day were less reflective of gISC on the other. This effect is greatly reduced in Fig. 2, suggesting that the unreliable low gISC values may reflect high levels of head motion. The scatterplots in this and subsequent supplementary figures show RSFC and ISC values before they were ranked and adjusted for head motion. As such, these best fit lines do not reflect the actual effects reported in the main text and are included here to facilitate visualization. (B) Visualization of the raw model predictions reveals that the model tended to overestimate gISC, especially for individuals with especially low Day 2 gISC. As rCPM was run on *z*-transformed gISC values, predicted gISC values were converted back to Pearson’s *r* for visualization in panel B.

**Figure S2.**
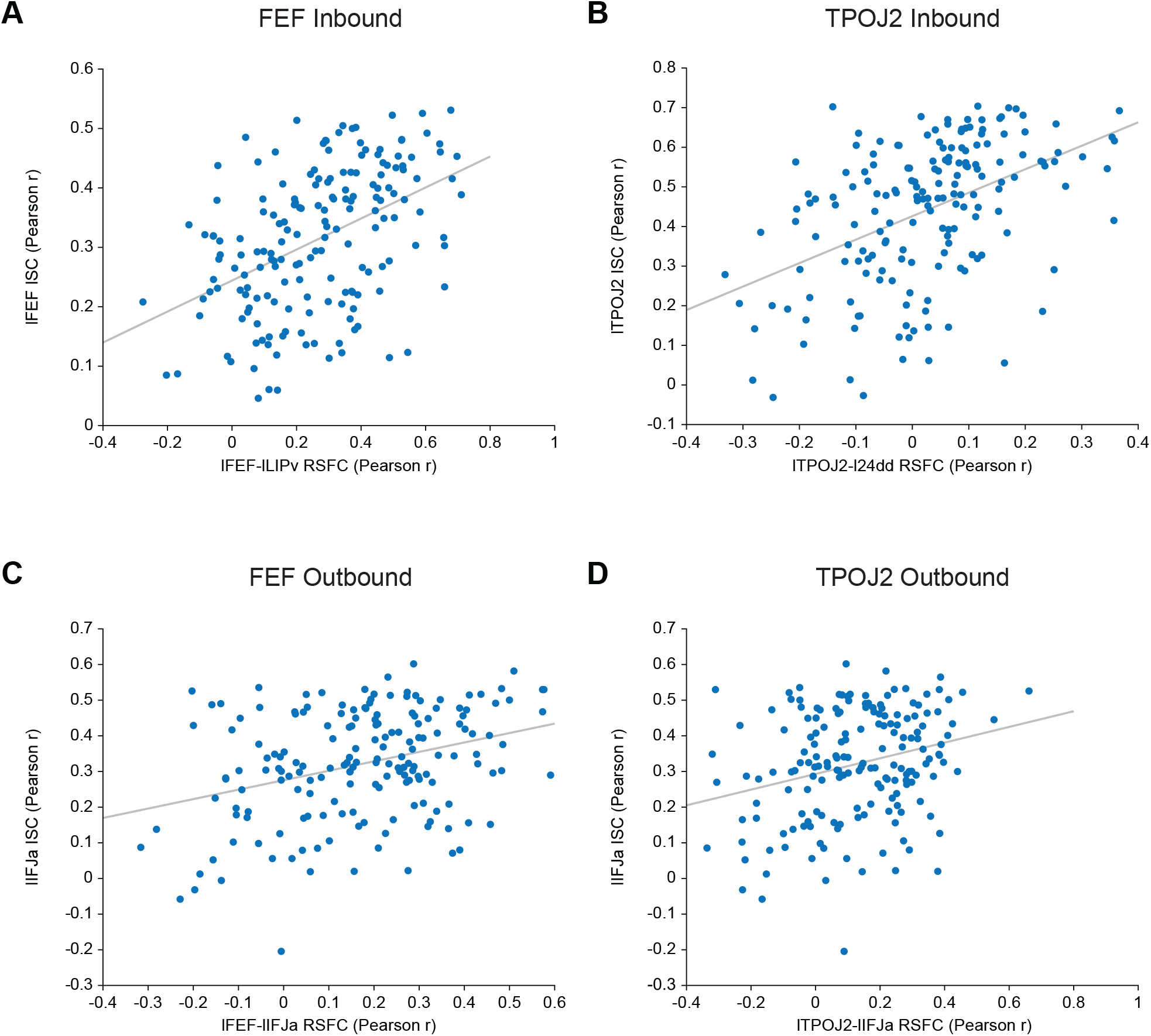
Raw inbound/outbound scatterplots. (A-D) These scatterplots reflect the raw ISC and RSFC inputs for the inbound/outbound relationships visualized in Fig. 3.

**Figure S3.**
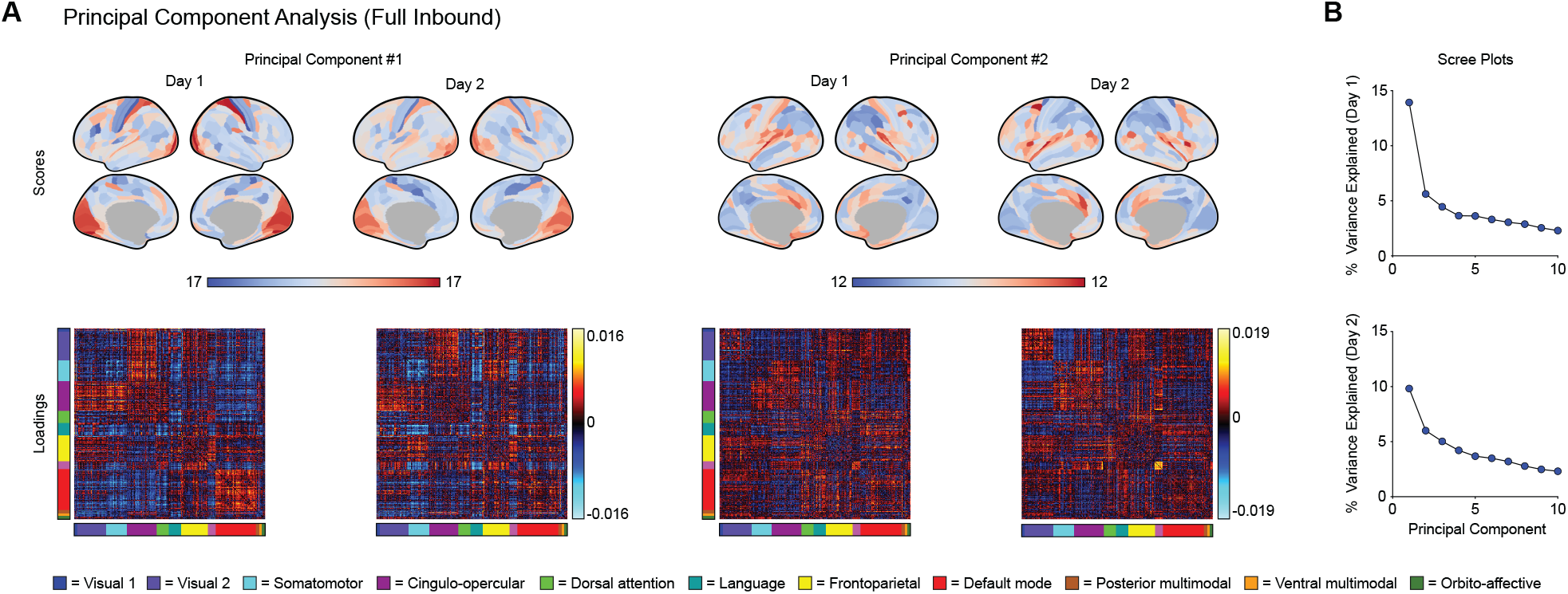
Principal component analysis of the full inbound matrices. (A) Cortical surfaces display the scores and RSFC heatmaps display the loadings for the first two principal components (PCs) of the full inbound matrices. For each PC, parcels that appear redder on the cortical surfaces exhibit greater ISC in individuals whose RSFC matrices look more like the loadings shown in the corresponding heatmaps. (B) Scree plots show that > 10 PCs are needed to account for a majority of the variance in the full inbound matrices, with the percentage of variance explained dropping off sharply after the first PC on both days.

**Figure S4.**
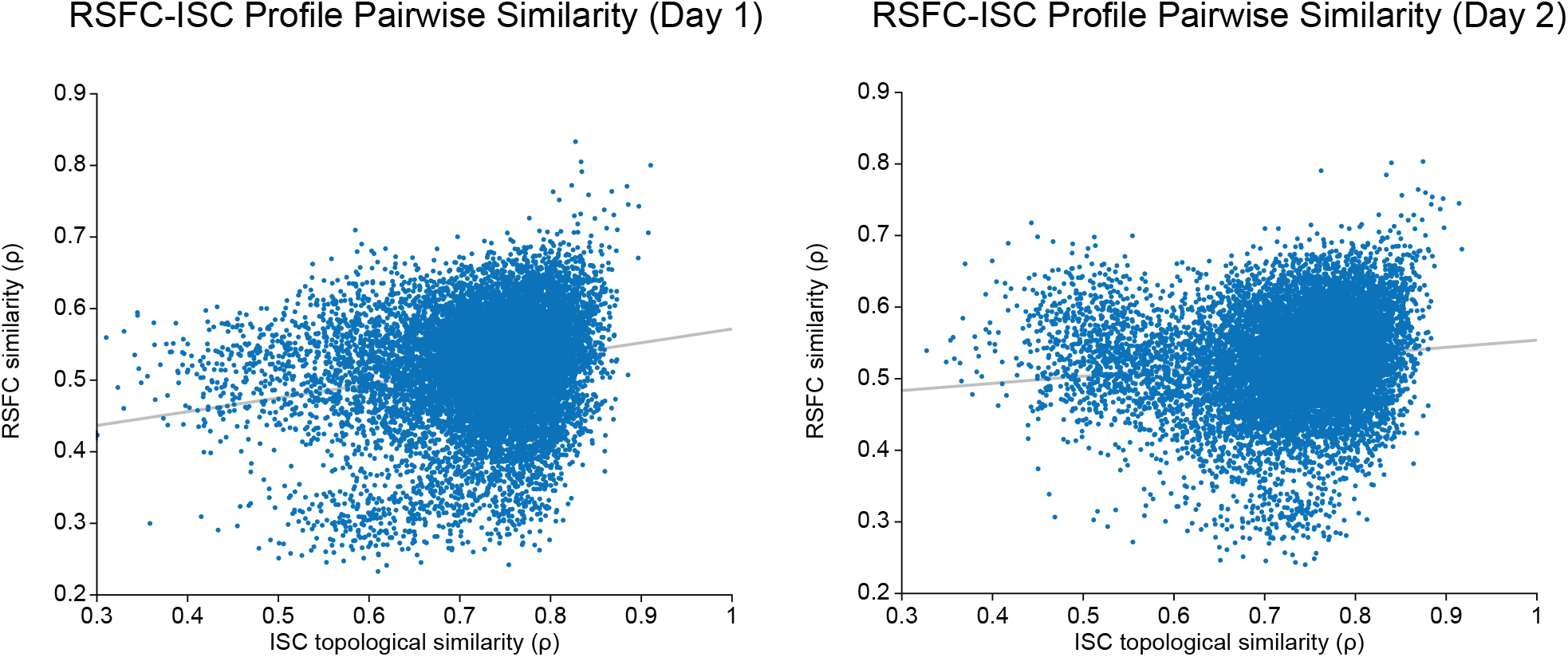
Raw RSFC-ISC similarity scatterplots. These scatterplots reflect the raw ISC and RSFC similarity inputs for the topological similarity relationships visualized in Fig. 6B.

## REFERENCES

1. Ambekar, A., Ward, C., Mohammed, J., Male, S., & Skiena, S. (2009). Name-ethnicity classification from open sources. Proceedings of the 15th ACM SIGKDD International Conference on Knowledge Discovery and Data Mining, 49–58. https://doi.org/10.1145/1557019.1557032

2. Arkin, S. C., Ruiz-Betancourt, D., Jamerson, E. C., Smith, R. T., Strauss, N. E., Klim, C. C., Javitt, D. C., & Patel, G. H. (2020). Deficits and compensation: Attentional control cortical networks in schizophrenia. NeuroImage. Clinical, 27, 102348. https://doi.org/10.1016/j.nicl.2020.102348

3. Baldassano, C., Chen, J., Zadbood, A., Pillow, J. W., Hasson, U., & Norman, K. A. (2017). Discovering event structure in continuous narrative perception and memory. Neuron, 95(3), 709–721.e5. https://doi.org/10.1016/j.neuron.2017.06.041

4. Campbell, K. L., Shafto, M. A., Wright, P., Tsvetanov, K. A., Geerligs, L., Cusack, R., Cam-CAN, & Tyler, L. K. (2015). Idiosyncratic responding during movie-watching predicted by age differences in attentional control. Neurobiology of Aging, 36(11), 3045–3055. https://doi.org/10.1016/j.neurobiolaging.2015.07.028

5. Caplar, N., Tacchella, S., & Birrer, S. (2017). Quantitative evaluation of gender bias in astronomical publications from citation counts. Nature Astronomy, 1(6), 0141. https://doi.org/10.1038/s41550-017-0141

6. Cole, M. W., Ito, T., Bassett, D. S., & Schultz, D. H. (2016). Activity flow over resting-state networks shapes cognitive task activations. Nature Neuroscience, 19(12), 1718–1726. https://doi.org/10.1038/nn.4406

7. Corbetta, M., Patel, G., & Shulman, G. L. (2008). The reorienting system of the human brain: from environment to theory of mind. Neuron, 58(3), 306–324. https://doi.org/10.1016/j.neuron.2008.04.017

8. Dion, M. L., Sumner, J. L., & Mitchell, S. M. (2018). Gendered citation patterns across political science and social science methodology fields. Political Analysis, 26(3), 312–327. https://doi.org/10.1017/pan.2018.12

9. Dosenbach, N. U. F., Visscher, K. M., Palmer, E. D., Miezin, F. M., Wenger, K. K., Kang, H. C., Burgund, E. D., Grimes, A. L., Schlaggar, B. L., & Petersen, S. E. (2006). A core system for the implementation of task sets. Neuron, 50(5), 799–812. https://doi.org/10.1016/j.neuron.2006.04.031

10. Dworkin, J. D., Linn, K. A., Teich, E. G., Zurn, P., Shinohara, R. T., & Bassett, D. S. (2020). The extent and drivers of gender imbalance in neuroscience reference lists. Nature Neuroscience, 23(8), 918–926. https://doi.org/10.1038/s41593-020-0658-y

11. Eickhoff, S. B., Milham, M., & Vanderwal, T. (2020). Towards clinical applications of movie fMRI. NeuroImage, 217, 116860. https://doi.org/10.1016/j.neuroimage.2020.116860

12. Elliott, M. L., Knodt, A. R., Ireland, D., Morris, M. L., Poulton, R., Ramrakha, S., Sison, M. L., Moffitt, T. E., Caspi, A., & Hariri, A. R. (2020). What is the test-retest reliability of common task-functional MRI measures? New empirical evidence and a meta-analysis. Psychological Science, 31(7), 792–806. https://doi.org/10.1177/0956797620916786

13. Fair, D. A., Miranda-Dominguez, O., Snyder, A. Z., Perrone, A., Earl, E. A., Van, A. N., Koller, J. M., Feczko, E., Tisdall, M. D., van der Kouwe, A., Klein, R. L., Mirro, A. E., Hampton, J. M., Adeyemo, B., Laumann, T. O., Gratton, C., Greene, D. J., Schlaggar, B. L., Hagler, D. J., Jr,… Dosenbach, N. U. F. (2020). Correction of respiratory artifacts in MRI head motion estimates. NeuroImage, 208, 116400. https://doi.org/10.1016/j.neuroimage.2019.116400

14. Felleman, D. J., & Van Essen, D. C. (1991). Distributed hierarchical processing in the primate cerebral cortex. Cerebral Cortex, 1(1), 1–47. https://doi.org/10.1093/cercor/1.1.1

15. Finn, E. S., & Bandettini, P. A. (2020). Movie-watching outperforms rest for functional connectivity-based prediction of behavior. bioRxiv. https://doi.org/10.1101/2020.08.23.263723

16. Finn, E. S., Corlett, P. R., Chen, G., Bandettini, P. A., & Constable, R. T. (2018). Trait paranoia shapes inter-subject synchrony in brain activity during an ambiguous social narrative. Nature Communications, 9(1), 2043. https://doi.org/10.1038/s41467-018-04387-2

17. Finn, E. S., Glerean, E., Khojandi, A. Y., Nielson, D., Molfese, P. J., Handwerker, D. A., & Bandettini, P. A. (2020). Idiosynchrony: From shared responses to individual differences during naturalistic neuroimaging. NeuroImage, 215, 116828. https://doi.org/10.1016/j.neuroimage.2020.116828

18. Finn, E. S., Shen, X., Scheinost, D., Rosenberg, M. D., Huang, J., Chun, M. M., Papademetris, X., & Constable, R. T. (2015). Functional connectome fingerprinting: identifying individuals using patterns of brain connectivity. Nature Neuroscience, 18(11), 1664–1671. https://doi.org/10.1038/nn.4135

19. Fox, M. D., Buckner, R. L., White, M. P., Greicius, M. D., & Pascual-Leone, A. (2012). Efficacy of transcranial magnetic stimulation targets for depression is related to intrinsic functional connectivity with the subgenual cingulate. Biological Psychiatry, 72(7), 595–603. https://doi.org/10.1016/j.biopsych.2012.04.028

20. Fox, M. D., Snyder, A. Z., McAvoy, M. P., Barch, D. M., & Raichle, M. E. (2005). The BOLD onset transient: identification of novel functional differences in schizophrenia. NeuroImage, 25(3), 771–782. https://doi.org/10.1016/j.neuroimage.2004.12.025

21. Gao, S., Greene, A. S., Constable, R. T., & Scheinost, D. (2019). Combining multiple connectomes improves predictive modeling of phenotypic measures. NeuroImage, 201, 116038. https://doi.org/10.1016/j.neuroimage.2019.116038

22. Glasser, M. F., Coalson, T. S., Robinson, E. C., Hacker, C. D., Harwell, J., Yacoub, E., Ugurbil, K., Andersson, J., Beckmann, C. F., Jenkinson, M., Smith, S. M., & Van Essen, D. C. (2016). A multi-modal parcellation of human cerebral cortex. Nature, 536(7615), 171–178. https://doi.org/10.1038/nature18933

23. Glasser, M. F., Sotiropoulos, S. N., Wilson, J. A., Coalson, T. S., Fischl, B., Andersson, J. L., Xu, J., Jbabdi, S., Webster, M., Polimeni, J. R., Van Essen, D. C., Jenkinson, M., & WU-Minn HCP Consortium. (2013). The minimal preprocessing pipelines for the Human Connectome Project. NeuroImage, 80, 105–124. https://doi.org/10.1016/j.neuroimage.2013.04.127

24. Gordon, E. M., Laumann, T. O., Gilmore, A. W., Newbold, D. J., Greene, D. J., Berg, J. J., Ortega, M., Hoyt-Drazen, C., Gratton, C., Sun, H., Hampton, J. M., Coalson, R. S., Nguyen, A. L., McDermott, K. B., Shimony, J. S., Snyder, A. Z., Schlaggar, B. L., Petersen, S. E., Nelson, S. M., & Dosenbach, N. U. F. (2017). Precision functional mapping of individual human brains. Neuron, 95(4), 791–807.e7. https://doi.org/10.1016/j.neuron.2017.07.011

25. Gratton, C., Dworetsky, A., Coalson, R. S., Adeyemo, B., Laumann, T. O., Wig, G. S., Kong, T. S., Gratton, G., Fabiani, M., Barch, D. M., Tranel, D., Miranda-Dominguez, O., Fair, D. A., Dosenbach, N. U. F., Snyder, A. Z., Perlmutter, J. S., Petersen, S. E., & Campbell, M. C. (2020). Removal of high frequency contamination from motion estimates in single-band fMRI saves data without biasing functional connectivity. NeuroImage, 217, 116866. https://doi.org/10.1016/j.neuroimage.2020.116866

26. Gratton, C., Laumann, T. O., Nielsen, A. N., Greene, D. J., Gordon, E. M., Gilmore, A. W., Nelson, S. M., Coalson, R. S., Snyder, A. Z., Schlaggar, B. L., Dosenbach, N. U. F., & Petersen, S. E. (2018). Functional brain networks are dominated by stable group and individual factors, not cognitive or daily variation. Neuron, 98(2), 439–452.e5. https://doi.org/10.1016/j.neuron.2018.03.035

27. Greene, A. S., Gao, S., Noble, S., Scheinost, D., & Constable, R. T. (2020). How tasks change whole-brain functional organization to reveal brain-phenotype relationships. Cell Reports, 32(8), 108066. https://doi.org/10.1016/j.celrep.2020.108066

28. Griffanti, L., Salimi-Khorshidi, G., Beckmann, C. F., Auerbach, E. J., Douaud, G., Sexton, C. E., Zsoldos, E., Ebmeier, K. P., Filippini, N., Mackay, C. E., Moeller, S., Xu, J., Yacoub, E., Baselli, G., Ugurbil, K., Miller, K. L., & Smith, S. M. (2014). ICA-based artefact removal and accelerated fMRI acquisition for improved resting state network imaging. NeuroImage, 95, 232–247. https://doi.org/10.1016/j.neuroimage.2014.03.034

29. Gruskin, D. C., Rosenberg, M. D., & Holmes, A. J. (2020). Relationships between depressive symptoms and brain responses during emotional movie viewing emerge in adolescence. NeuroImage, 216, 116217. https://doi.org/10.1016/j.neuroimage.2019.116217

30. Guo, C. C., Nguyen, V. T., Hyett, M. P., Parker, G. B., & Breakspear, M. J. (2015). Out-of-sync: disrupted neural activity in emotional circuitry during film viewing in melancholic depression. Scientific Reports, 5, 11605. https://doi.org/10.1038/srep11605

31. Hasson, U., Nir, Y., Levy, I., Fuhrmann, G., & Malach, R. (2004). Intersubject synchronization of cortical activity during natural vision. Science, 303(5664), 1634–1640. https://doi.org/10.1126/science.1089506

32. Ito, T., Kulkarni, K. R., Schultz, D. H., Mill, R. D., Chen, R. H., Solomyak, L. I., & Cole, M. W. (2017). Cognitive task information is transferred between brain regions via resting-state network topology. Nature Communications, 8(1), 1027. https://doi.org/10.1038/s41467-017-01000-w

33. Ji, J. L., Spronk, M., Kulkarni, K., Repovš, G., Anticevic, A., & Cole, M. W. (2019). Mapping the human brain’s cortical-subcortical functional network organization. NeuroImage, 185, 35–57. https://doi.org/10.1016/j.neuroimage.2018.10.006

34. Ki, J. J., Kelly, S. P., & Parra, L. C. (2016). Attention strongly modulates reliability of neural responses to naturalistic narrative stimuli. The Journal of Neuroscience, 36(10), 3092–3101. https://doi.org/10.1523/JNEUROSCI.2942-15.2016

35. Kragel, P., Han, X., Kraynak, T., Gianaros, P. J., & Wager, T. (2020). fMRI can be highly reliable, but it depends on what you measure. PsyArXiv. https://doi.org/10.31234/osf.io/9eaxk

36. Lerner, Y., Honey, C. J., Silbert, L. J., & Hasson, U. (2011). Topographic mapping of a hierarchy of temporal receptive windows using a narrated story. The Journal of Neuroscience, 31(8), 2906–2915. https://doi.org/10.1523/JNEUROSCI.3684-10.2011

37. Li, J., Kong, R., Liégeois, R., Orban, C., Tan, Y., Sun, N., Holmes, A. J., Sabuncu, M. R., Ge, T., & Yeo, B. T. T. (2019). Global signal regression strengthens association between resting-state functional connectivity and behavior. NeuroImage, 196, 126–141. https://doi.org/10.1016/j.neuroimage.2019.04.016

38. Maliniak, D., Powers, R., & Walter, B. F. (2013). The gender citation gap in international relations. International Organization, 67(4), 889–922. https://doi.org/10.1017/S0020818313000209

39. Marcus, D. S., Harwell, J., Olsen, T., Hodge, M., Glasser, M. F., Prior, F., Jenkinson, M., Laumann, T., Curtiss, S. W., & Van Essen, D. C. (2011). Informatics and data mining tools and strategies for the human connectome project. Frontiers in Neuroinformatics, 5, 4. https://doi.org/10.3389/fninf.2011.00004

40. Mesulam, M. M. (1998). From sensation to cognition. Brain, 121 (Pt 6), 1013–1052. https://doi.org/10.1093/brain/121.6.1013

41. Mill, R. D., Gordon, B. A., Balota, D. A., & Cole, M. W. (2020). Predicting dysfunctional age-related task activations from resting-state network alterations. NeuroImage, 221, 117167. https://doi.org/10.1016/j.neuroimage.2020.117167

42. Mitchell, S. M., Lange, S., & Brus, H. (2013). Gendered citation patterns in international relations journals. International Studies Perspectives, 14(4), 485–492. https://doi.org/10.1111/insp.12026

43. Nastase, S. A., Gazzola, V., Hasson, U., & Keysers, C. (2019). Measuring shared responses across subjects using intersubject correlation. Social Cognitive and Affective Neuroscience, 14(6), 667–685. https://doi.org/10.1093/scan/nsz037

44. O’Connor, D., Lake, E. M. R., Scheinost, D., & Constable, R. T. (2020). Bootstrap aggregating improves the generalizability of Connectome Predictive Modelling. bioRxiv. https://doi.org/10.1101/2020.07.08.193664

45. Pajula, J., & Tohka, J. (2016). How many is enough? Effect of sample size in inter-subject correlation analysis of fMRI. Computational Intelligence and Neuroscience, 2016. https://doi.org/10.1155/2016/2094601

46. Patel, G. H., Sestieri, C., & Corbetta, M. (2019). The evolution of the temporoparietal junction and posterior superior temporal sulcus. Cortex, 118, 38–50. https://doi.org/10.1016/j.cortex.2019.01.026

47. Paus, T. (1996). Location and function of the human frontal eye-field: A selective review. Neuropsychologia, 34(6), 475–483. https://doi.org/10.1016/0028-3932(95)00134-4

48. Regev, M., Simony, E., Lee, K., Tan, K. M., Chen, J., & Hasson, U. (2019). Propagation of information along the cortical hierarchy as a function of attention while reading and listening to stories. Cerebral Cortex, 29(10), 4017–4034. https://doi.org/10.1093/cercor/bhy282

49. Rosenberg, M. D., Finn, E. S., Scheinost, D., Papademetris, X., Shen, X., Constable, R. T., & Chun, M. M. (2016). A neuromarker of sustained attention from whole-brain functional connectivity. Nature Neuroscience, 19(1), 165–171. https://doi.org/10.1038/nn.4179

50. Rosenberg, M. D., Scheinost, D., Greene, A. S., Avery, E. W., Kwon, Y. H., Finn, E. S., Ramani, R., Qiu, M., Constable, R. T., & Chun, M. M. (2020). Functional connectivity predicts changes in attention observed across minutes, days, and months. Proceedings of the National Academy of Sciences of the United States of America, 117(7), 3797–3807. https://doi.org/10.1073/pnas.1912226117

51. Salehi, M., Greene, A. S., Karbasi, A., Shen, X., Scheinost, D., & Constable, R. T. (2020). There is no single functional atlas even for a single individual: Functional parcel definitions change with task. NeuroImage, 208, 116366. https://doi.org/10.1016/j.neuroimage.2019.116366

52. Salmi, J., Roine, U., Glerean, E., Lahnakoski, J., Nieminen-von Wendt, T., Tani, P., Leppämäki, S., Nummenmaa, L., Jääskeläinen, I. P., Carlson, S., Rintahaka, P., & Sams, M. (2013). The brains of high functioning autistic individuals do not synchronize with those of others. NeuroImage. Clinical, 3, 489–497. https://doi.org/10.1016/j.nicl.2013.10.011

53. Scheinost, D., Noble, S., Horien, C., Greene, A. S., Lake, E. M., Salehi, M., Gao, S., Shen, X., O’Connor, D., Barron, D. S., Yip, S. W., Rosenberg, M. D., & Constable, R. T. (2019). Ten simple rules for predictive modeling of individual differences in neuroimaging. NeuroImage, 193, 35–45. https://doi.org/10.1016/j.neuroimage.2019.02.057

54. Shen, X., Finn, E. S., Scheinost, D., Rosenberg, M. D., Chun, M. M., Papademetris, X., & Constable, R. T. (2017). Using connectome-based predictive modeling to predict individual behavior from brain connectivity. Nature Protocols, 12(3), 506–518. https://doi.org/10.1038/nprot.2016.178

55. Song, H., Finn, E. S., & Rosenberg, M. D. (2020). Neural signatures of attentional engagement during narratives and its consequences for event memory. bioRxiv. https://doi.org/10.1101/2020.08.26.266320

56. Sood, G., & Laohaprapanon, S. (2018). Predicting race and ethnicity from the sequence of characters in a name. arXiv. http://arxiv.org/abs/1805.02109

57. Tavor, I., Parker Jones, O., Mars, R. B., Smith, S. M., Behrens, T. E., & Jbabdi, S. (2016). Task-free MRI predicts individual differences in brain activity during task performance. Science, 352(6282), 216–220. https://doi.org/10.1126/science.aad8127

58. van den Heuvel, M. P., & Hulshoff Pol, H. E. (2010). Exploring the brain network: A review on resting-state fMRI functional connectivity. European Neuropsychopharmacology, 20(8), 519–534. https://doi.org/10.1016/j.euroneuro.2010.03.008

59. Van Essen, D. C., Smith, S. M., Barch, D. M., Behrens, T. E. J., Yacoub, E., Ugurbil, K., & WU-Minn HCP Consortium. (2013). The WU-Minn Human Connectome Project: an overview. NeuroImage, 80, 62–79. https://doi.org/10.1016/j.neuroimage.2013.05.041

60. Váša, F., Seidlitz, J., Romero-Garcia, R., Whitaker, K. J., Rosenthal, G., Vértes, P. E., Shinn, M., Alexander-Bloch, A., Fonagy, P., Dolan, R. J., Jones, P. B., Goodyer, I. M., NSPN consortium, Sporns, O., & Bullmore, E. T. (2018). Adolescent tuning of association cortex in human structural brain networks. Cerebral Cortex, 28(1), 281–294. https://doi.org/10.1093/cercor/bhx249

61. Weinstein, S. M., Vandekar, S. N., Adebimpe, A., Tapera, T. M., Robert-Fizgerald, T., Gur, R. C., Gur, R. E., Raznahan, A., Satterthwaite, T. D., Alexander-Bloch, A. F., & Shinohara, R. T. (2020). A simple permutation-based test of intermodal correspondence. bioRxiv. https://doi.org/10.1101/2020.09.10.285049

62. Wig, G. S. (2017). Segregated systems of human brain networks. Trends in Cognitive Sciences, 21(12), 981–996. https://doi.org/10.1016/j.tics.2017.09.006

63. Zhou, D., Stiso, J., Cornblath, E., Teich, E., Blevins, A. S., Oudyk, K., Michael, C. Virtualmario, & Camp, C. (2020). dalejn/cleanBib: v1.1.1. https://doi.org/10.5281/zenodo.4104748

